# Localization of the pioneer factor GAF to subnuclear foci is driven by DNA binding and required to silence satellite repeat expression

**DOI:** 10.1101/2022.11.29.518380

**Authors:** Marissa. M. Gaskill, Isabella V. Soluri, Annemarie E. Branks, Alan P. Boka, Michael R. Stadler, Katherine Vietor, Hao-Yu S. Huang, Tyler J. Gibson, Mustafa Mir, Shelby A. Blythe, Melissa M. Harrison

## Abstract

The eukaryotic genome is organized to enable the precise regulation of gene expression required for development. This organization is established during early development when the embryo transitions from a fertilized germ cell to the totipotent zygote. To understand the factors and processes that drive genomic organization, we focused on the pioneer factor GAGA factor (GAF) that is required for early embryonic development in *Drosophila.* GAF transcriptionally activates the zygotic genome and is localized to subnuclear foci. We show that this non-uniform distribution is driven by binding to the highly abundant GA-satellite repeats. At GA-repeats, GAF is necessary to form heterochromatin and silence transcription. Thus, GAF is required to establish both active and silent regions. We propose that foci formation enables GAF to have opposing transcriptional roles within a single nucleus. Our data support a model in which modulation of the subnuclear concentration of transcription factors acts to organize the nucleus into functionally distinct domains that are essential for the robust regulation of gene expression.

## Introduction

Development requires the precise control of gene expression. Transitions in cell fate necessitate both the activation and silencing of transcription controlled by *trans*-acting factors that bind chromatin. However, chromatin presents a barrier to the binding of many transcription factors (Li et al. 2011; Zhu et al. 2018). By contrast, pioneer transcription factors can bind to sequence-specific target motifs when they are wrapped around a nucleosome (Zaret 2020; Larson et al. 2021; Balsalobre and Drouin 2022). These factors act at the top of gene regulatory networks to drive widespread changes in cell fate by creating local regions of chromatin accessibility that can be bound by additional transcription factors (Larson et al. 2021; Balsalobre and Drouin 2022). Silencing of gene expression is associated with a loss of chromatin accessibility at *cis*-regulatory regions and the deposition of histone modifications that inhibit transcription-factor binding (Allshire and Madhani 2018). Gene expression programs controlled by *trans-*acting factors must be carefully balanced to achieve the changes in cell fate required during development (Larson et al. 2021; Allshire and Madhani 2018). Nonetheless, much remains unknown about how this process is regulated to precisely control cell-fate changes.

Dramatic changes in cell fate and gene expression occur during the rapid reprogramming of the fertilized egg to the totipotent embryo. Initially following fertilization, the zygotic genome is transcriptionally quiescent, and development is controlled by maternally deposited mRNAs and proteins. The genome is devoid of most histone modifications and lacks features of mature heterochromatin domains. Thus, transcriptional activation and silencing need to be established *de novo* in the embryo during this maternal-to-zygotic transition (MZT) (Schulz and Harrison 2019; Vastenhouw et al. 2019). Initially following fertilization, chromatin in the nucleus largely lacks three-dimensional structure. As the transcriptional program is established, active and silent genomic regions are segregated into distinct compartments, and some *trans*-acting factors become localized to discrete subnuclear domains (Schulz and Harrison 2019; Vastenhouw et al. 2019; Zhang and Xie 2022; Strom et al. 2017; Hur et al. 2020). Together these events restructure the genome and lead to the rapid reprogramming of cell fate.

Several regulatory proteins and transcription factors have been reported to have non-uniform distribution within the nucleus, visualized as foci and referred to as condensates or hubs. In many cases, this distribution is driven by multivalent interactions between the intrinsically disordered regions (IDRs) prevalent in eukaryotic transcription factors (Boeynaems et al. 2018; Rippe 2022; Peng et al. 2020; Ferrie et al. 2022). For example, Heterochromatin Protein 1a/*α* (HP1a/*α*) forms membraneless condensates thought to be formed by liquid-liquid phase separation (LLPS) in both flies and mammals (Strom et al. 2017; Larson et al. 2017). In contrast to the stable condensates formed by HP1, the *Drosophila* transcription factor Zelda forms dynamic transcription factor hubs that mediate DNA binding of additional transcription factors (Mir et al. 2017, 2018b; Dufourt et al. 2018; Yamada et al. 2019). Functionally similar, high-local concentration microenvironments have also been reported for the transcription factor Ultrabithorax (Tsai et al. 2017). These transcription factor condensates have been proposed to be important for a number of processes, including active transcription (Boija et al. 2018). Such condensates or hubs are suggested to be functionally significant in partitioning the genome and determining transcriptional activity. However, it has been challenging to test the functional significance of non-uniform subnuclear distribution of proteins within a biological system (Alberti et al. 2019).

We employed the rapidly developing, highly genetically tractable model organism *Drosophila melanogaster* to study the impact of subnuclear domains during the dynamic reprogramming in the early embryo. Early *Drosophila* development is characterized by 13 synchronous nuclear divisions, which alternate between replication and mitosis with no gap phases. At nuclear cycle (NC) 8 transcription from some of the earliest zygotic genes can be detected, but widespread zygotic genome activation (ZGA) does not occur until NC14 (Hamm and Harrison 2018). We recently showed that the pioneering factor, GAGA factor (GAF) is required for development through the MZT. GAF activates the zygotic genome, preferentially driving gene expression during the major wave of ZGA at NC14 (Gaskill et al. 2021). During early development, GAF forms subnuclear foci that are retained on chromatin during mitosis (Raff et al. 1994; Gaskill et al. 2021). While these GAF foci were first observed decades ago, their contribution to transcriptional regulation remains unknown.

We used GAF foci as a model to understand protein features that drive non-uniform distribution of pioneer factors and how this distribution contributes to transcription-factor function during a dynamic reprogramming period. Unexpectedly, we determined that GAF localization to foci is driven by DNA-binding activity rather than IDR-mediated interactions. GAF foci are regions of high local GAF concentration directed by DNA binding at GA-rich satellite repeats. GAF binding at these repeats is essential for H3K9me3 deposition and transcriptional silencing. Thus, at these regions GAF functions as a transcriptional repressor and contributes to the establishment of mature heterochromatin. Based on this work and our previous data, we conclude that GAF simultaneously functions to reprogram both the active and the silent genome during the MZT in *Drosophila.* In high concentration foci, GAF is necessary to establish a transcriptionally silent state, while the more diffuse populations of GAF activate transcription.

## Results

### GAF forms foci during the MZT

GAGA factor, like several other transcription factors, is non-uniformly distributed in the nucleus (Raff et al. 1994). Imaging endogenous GAF tagged with super-folder GFP (sfGFP), we demonstrated that GAF forms robust, discrete foci during interphase of early embryogenesis and is mitotically retained (Gaskill et al. 2021) (Figure 1A). Using high resolution lattice light-sheet imaging on sfGFP-tagged GAF embryos through NC13 and NC14, we identified that during NC13 GAF forms on average 8 foci per nucleus. (Chen et al. 2014). During NC13, as the chromatin compacts at the entry into mitosis, the foci decreased in number and volume (Figure S1A-D). In both NC13 and NC14, shifts in the percent of total nuclear volume occupied by GAF foci over time mirrored the changes in total foci volume (Figure S1A). The similarity in these profiles indicated that differences in total foci volume were not simply due to fluctuations in nuclear volume. A similar number of GAF foci were present at the start of NC14 (∼5 per nucleus) as were observed during NC13 (Figure 1B, S1C). However, at approximately 17 minutes into interphase of NC14 the number and volume of foci began to decrease (Figure 1B, 1C, S1A). Unlike during NC13 when foci number decreased upon mitotic entry, the decrease in foci number and volume during NC14 did not coincide with a global reorganization of foci or entry into mitosis, suggesting that this decrease might be caused by fusion of individual foci (Figure S1D). Indeed, we observed fusion of GAF foci at NC14 (Figure 1D). The localization of GAF to concentrated foci that undergo fusion is reminiscent of phase-separated nuclear factors (Strom et al. 2017; Hur et al. 2020), or membraneless compartments formed by multivalent interactions between intrinsically disordered regions (IDRs) (Boeynaems et al. 2018; Rippe 2022; Peng et al. 2020). GAF has an intrinsically disordered poly-glutamine (poly-Q) enriched C-terminal domain, which drives multimerization *in vitro* (Wilkins and Lis 1999). Thus, we sought to investigate the features that caused the localization of GAF to discrete subnuclear foci and whether this localization was driven by intrinsically disordered regions (IDRs) as has been reported for other factors.

**Figure 1:**
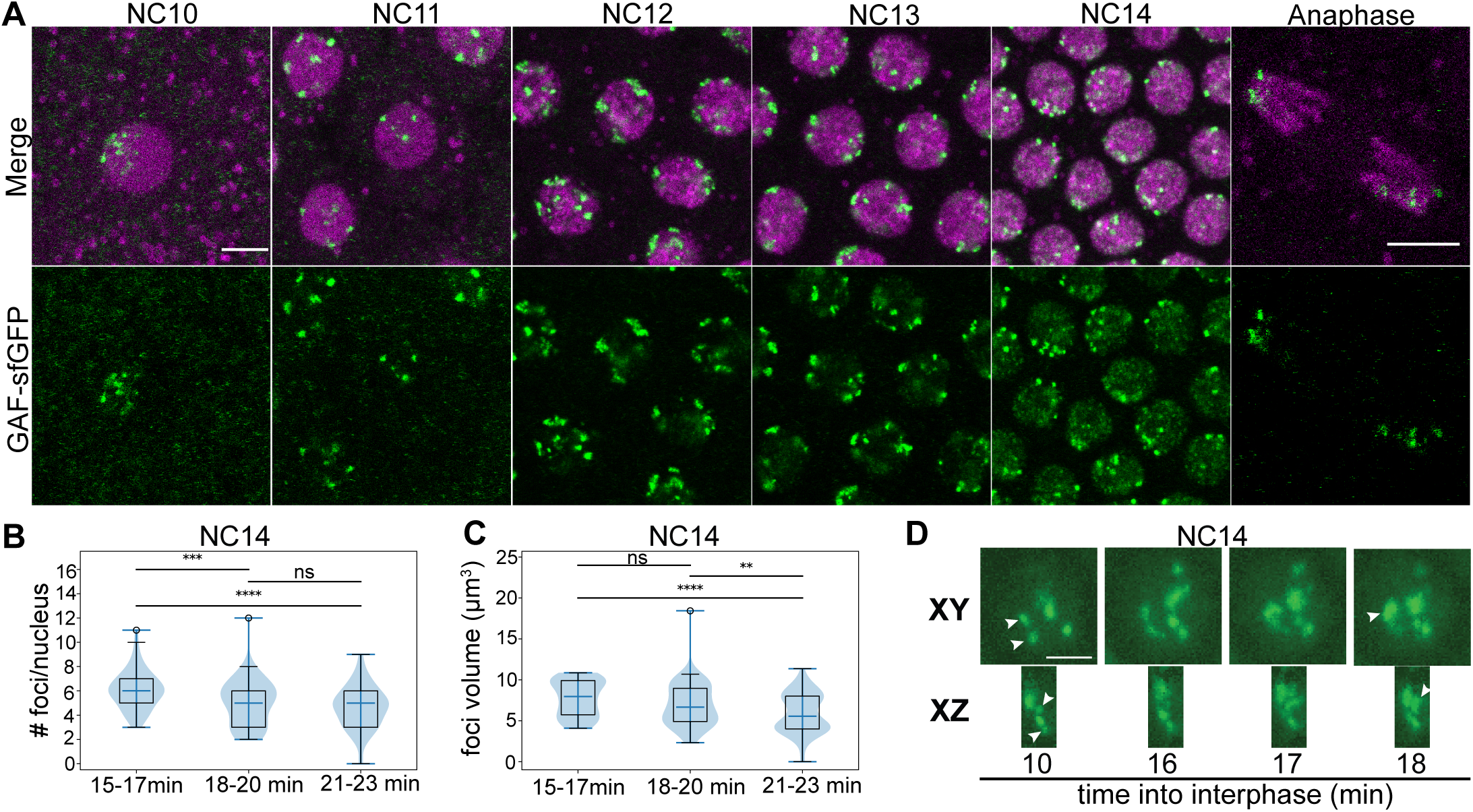
GAF forms multiple, stable nuclear foci during the MZT. A. Representative images of embryos from NC10-14 (as indicated) laid by *His2Av-RFP; GAF-sfGFP* females. GAF-sfGFP is in green. His2Av-RFP is in magenta. Scale bars, 5μM. B. Quantification of the number of sfGFP tagged GAF foci per nucleus in NC14. C. Quantification of the volume of sfGFP tagged GAF in NC14. Asterisks indicate pairwise p-value thresholds. ** = 0.01, *** = 0.001, **** = 0.0001 (Tukey-Kramer test). n = 2 embryos, 29 nuclei analyzed. D. Orthogonal x-y and x-z views of sfGFP tagged GAF foci fusion during NC14. Scale bar, 2.5μm.

### GAF isoforms have overlapping functions

There are two predominant isoforms of GAF that differ in the length and sequence of their C-terminal poly-Q domains: a long isoform (582 aa) and a short isoform (519 aa) (Figure S2A). If the poly-Q IDR promotes localization to foci, it is possible that the isoform-specific domains promote distinct abilities of each isoform to localize to subnuclear foci and discrete *in vivo* functions. The GAF isoforms have developmentally distinct expression patterns (Benyajati et al. 1997). Analysis of modENCODE RNA-seq data demonstrated that the transcript for the long isoform is not present in the 0–2-hour embryo (Figure S2B). Furthermore, the protein encoded by this isoform is not detectable in the embryo at this time in development (Gaskill et al. 2021; Benyajati et al. 1997). This distinct developmental expression pattern and the high level of conservation of the two GAF isoforms in distantly related *Drosophila* species suggests that the two isoforms may have separate *in vivo* functional roles (Lintermann et al. 1998). By contrast, prior studies using transgene expression concluded that the two isoforms largely overlap in their function (Greenberg and Schedl 2001). Because transgenes do not always reflect endogenous expression levels and patterns, we made mutations in the endogenous GAF locus using Cas9-gene editing to establish the functional roles of each isoform and to determine if the different IDR lengths were functionally relevant.

To interfere with the long isoform, a stop codon was introduced at the beginning of the 6^th^ exon, which is a coding region only in the long isoform (Figure 2A). This allele is referred to as *GAF^L^*. To interfere with the short isoform, the 297 bp coding region unique to the short isoform, including the stop codon was deleted (424-519aa). This region is alternatively spliced out of the long isoform transcript (Figure 2A). This results in the deletion of the C-terminal poly-Q domain from the short isoform and is referred to as *GAF^SΔPQ^*. Using reverse-transcriptase PCR and sequencing, we demonstrated that *GAF^L^* does not produce a detectable product for the long isoform, which is likely degraded by nonsense mediated decay (Figure 2A, S2B, S2C). Thus, flies homozygous for *GAF^L^* lack the long isoform. *GAF^SΔPQ^* results in a truncated stable transcript of the short isoform (Figure S2B-D), and immunoblot confirmed the existence of a truncated stable protein (Figure S2E). The long isoform was not detectable in these blots, likely because it is expressed at much lower levels than the short isoform (Gaskill et al. 2021). Thus, flies homozygous for *GAF^SΔPQ^* express the long isoform and an altered version of the short GAF protein lacking the entirety of the poly-Q domain.

**Figure 2:**
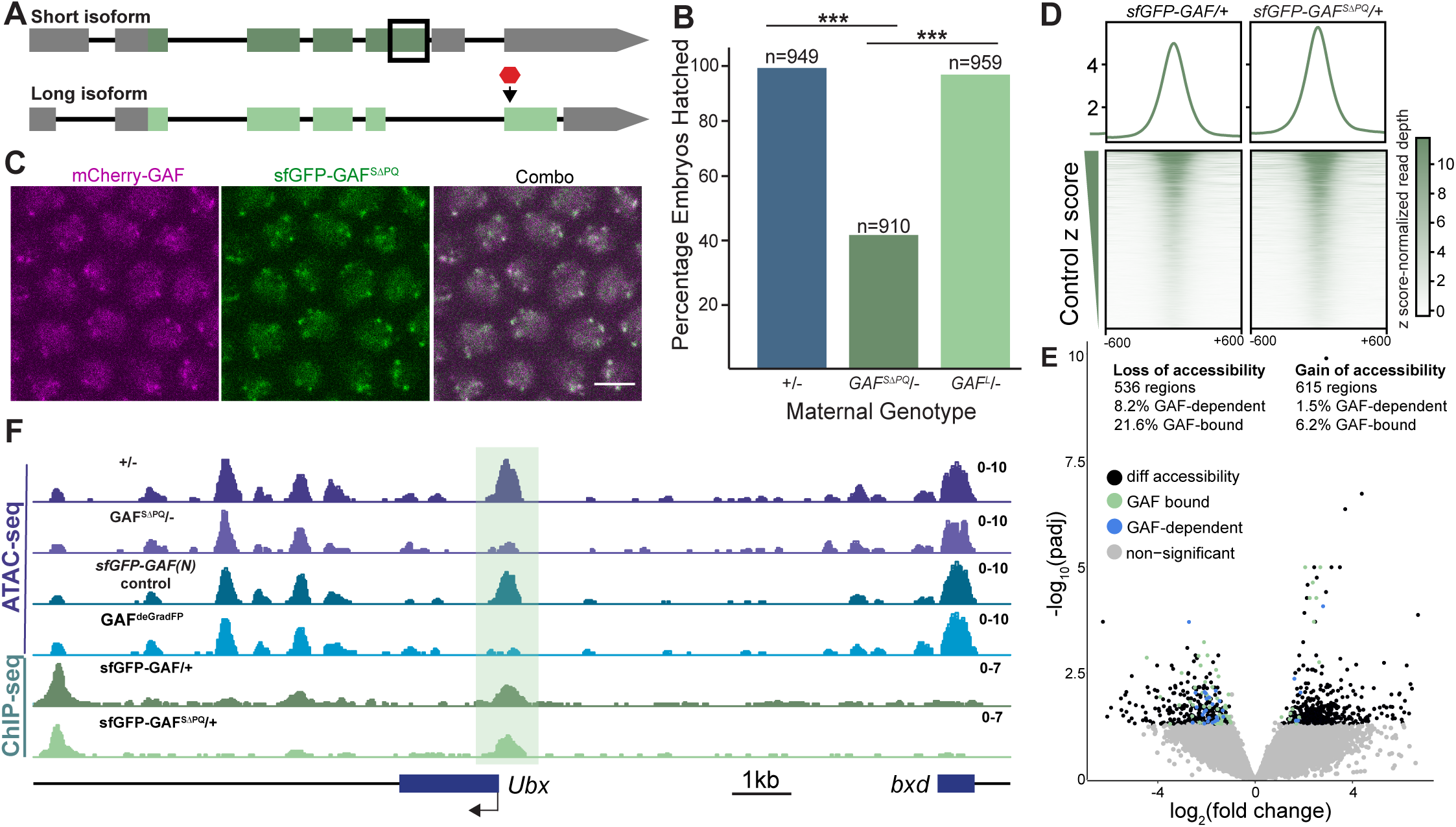
The intrinsically disordered poly-Q domain is not required for foci formation but contributes to pioneering activity. A. Model of two GAF splice isoforms. Coding regions are shown in green. Untranslated regions (UTRs) are in grey. Black lines indicate introns. Isoform specific mutations generated in this study are indicated. The black box denotes the region deleted in the short-isoform specific deletion. The red octagon indicates the location of the stop codon introduced in the long isoform. B. Percentage of hatched embryos laid by the maternal genotypes indicated crossed to *w^1118^* males (***, *χ*^2^, p = 2.2 x 10^-16^) n = total number of embryos assayed. C. Representative images of interphase NC14 embryos laid by *mCherry-GAF/sfGFP-GAF^SΔPQ^* females. mCherry-GAF is in magenta. sfGFP-GAF^SΔPQ^ is in green. Scale bars, 5μM. D. Heatmaps of anti-GFP ChIP-seq from 2-2.5hr AEL embryos laid by *sfGFP-GAF^SΔPQ^/+* females and control *sfGFP-GAF/+* females. Heatmaps are ordered by z score-normalized signal from control embryos. E. Volcano plot of regions that change in accessibility in 2-2.5hr AEL embryos laid by *GAF^SΔPQ^/-* females compared to embryos laid by *+/-* females. Stage 5 sfGFP-GAF(N) ChIP-seq (Gaskill et al. 2021) was used to identify GAF-bound regions, and GAF-dependent regions are those identified in Gaskill et al. 2021 as changing in accessibility in GAF^deGradFP^ embryos compared to controls. F. Genome browser tracks of ATAC-seq on 2-2.5hr AEL *sfGFP-GAF(N)* homozygous and GAF^deGradFP^ embryos (Gaskill et al. 2021) and 2-2.5hr AEL embryos laid by *GAF^SΔPQ^/-* females and control *+/-* females. Binding is indicated by anti-GFP ChIP-seq from 2-2.5hr AEL embryos laid by *sfGFP-GAF^SΔPQ^/+* and *sfGFP-GAF/+* females. All ATAC-seq and ChIP-seq genome browser tracks are z score-normalized. Region highlighted in green indicates the GAF-dependent, polyQ-domain sensitive, GAF-bound *Ubx* promoter.

Having generated isoform-specific alleles, we tested whether these alleles resulted in observable mutant phenotypes. GAF is essential for viability, and flies lacking zygotically expressed GAF die before the third instar larval stage (Farkas et al. 1994). Hypomorphic GAF alleles result in homeotic transformations (Farkas et al. 1994; Greenberg and Schedl 2001). By contrast, males and females with the *GAF^L^* or *GAF^SΔPQ^* allele over a GAF null allele were viable and fertile without any obvious mutant phenotypes. When we quantified adult viability for each allele in *trans* to a null allele, we confirmed that flies homozygous for the null allele failed to reach adulthood (Figure S2F). However, we identified no decrease in adult viability for either of the isoform-specific alleles (Figure S2F). Thus, the long isoform and the poly-Q domain of the short isoform are not individually required for adult viability.

### The GAF poly-Q domain is not required for foci localization, but contributes to pioneering activity

Given the previously reported roles of the poly-Q domain in transcriptional activation, protein multimerization and DNA distortion *in vitro* (Wilkins and Lis 1999; Vaquero et al. 2000; Agianian et al. 1999), we were surprised that flies lacking this domain in the short isoform showed no defects in adult viability. To further investigate the functional relevance of this domain, we examined the first few hours of embryonic development when only the short isoform is present. At this time, development is controlled by maternally provided products, and we recently demonstrated that GAF is essential (Gaskill et al. 2021). Thus, we tested the viability of embryos that inherited only the short isoform lacking the poly-Q domain from their mothers. Females heterozygous for the GAF null allele and either *GAF^L^*, *GAF^SΔPQ^*, or wild-type GAF were mated to *w^1118^* males. While 94% of embryos inheriting mRNA encoding wild-type GAF hatched, only 41.6% of embryos inheriting the poly-Q deletion mutant of the short isoform hatched (*χ*^2^, p= 2.2 x 10^-16^) (Figure 2B). Embryos inheriting only the short isoform (*GAF^L^*) had a 92.1% hatching rate, similar to the wild-type control (Figure 2B). This suggests that the poly-Q domain supports, but is not absolutely required, for GAF function.

IDRs can drive protein aggregation and phase transitions, and the poly-Q domain of GAF has been implicated in such functions (Boeynaems et al. 2018; Wilkins and Lis 1999; Agianian et al. 1999; Ferrie et al. 2022). Thus, it is possible that the reduced hatching rate for embryos inheriting the short isoform lacking the poly-Q domain might result from a failure of the mutant GAF protein to concentrate in subnuclear foci. To identify the localization pattern of GAF lacking the poly-Q domain, we engineered the endogenous 297bp short isoform poly-Q deletion in the background of a line in which we previously tagged endogenous GAF with super-folder GFP (sfGFP) to create *sfGFP-GAF^SΔPQ^* (Figure S2G) (Gaskill et al. 2021). In embryos inheriting this allele from their mothers, sfGFP-GAF^SΔPQ^ continued to localize to discrete foci, and these foci completely overlapped with full-length GAF endogenously tagged with mCherry at NC14 (Figure 2C). Therefore, the poly-Q IDR is dispensable for the recruitment of GAF to subnuclear foci. We performed chromatin immunoprecipitation coupled with high-throughput sequencing (ChIP-seq) on stage 5 embryos inheriting a single copy of maternally deposited *sfGFP-GAF^SΔPQ^* to test whether the deletion of the poly-Q domain affected genomic occupancy (Table S1). Spike-in normalized ChIP-seq data demonstrated that sfGFP-GAF^SΔPQ^ bound to 88% (2708 out of 3078) of the regions bound by sfGFP-GAF in the control embryos (Figure 2D). Together these data reveal that contrary to the expected role of the intrinsically disordered poly-Q domain in protein binding and localization, this domain is not required for recruitment to subnuclear foci or for chromatin binding.

Recently, it was shown in hemocytes that the GAF poly-Q domain contributes to stable chromatin binding (Tang et al. 2022). Compared to other transcription factors, GAF has a high residence time on chromatin (∼10 s versus ∼100 s), and it was proposed that this stable occupancy promotes chromatin accessibility and may be important for pioneering activity (Tang et al. 2022). We previously defined a role for GAF as a pioneer factor important for chromatin accessibility at a subset of genomic regions at NC14 (Gaskill et al. 2021). To directly test if the poly-Q domain contributes to GAF pioneering activity during the MZT, we used ATAC-seq to assay chromatin accessibility in NC14 embryos inheriting only GAF^SΔPQ^ from their mothers. When compared to controls, we identified hundreds of regions that changed in accessibility (Figures 2E, Table S2). The regions that lose accessibility were significantly enriched for GAF-binding sites when compared to unchanged accessible regions, with over 20% of sites that lose accessibility directly bound by GAF at stage 5 (Gaskill et al. 2021) (p-value < 2.2x10^-16^, log2(odds ratio)=1.5, two-tailed Fisher’s exact test). For example, at the *Ultrabithorax* (*Ubx*) promoter, GAF lacking the poly-Q domain was bound to chromatin, but binding was not sufficient for chromatin accessibility (Figure 2F). This contrasts with wild-type GAF function at the *Ubx* promoter, where GAF can bind to closed chromatin and drive accessibility (Gaskill et al., 2021). Together these data demonstrate that GAF lacking the poly-Q domain retains the ability to bind chromatin but is not sufficient to establish accessibility at a subset of these regions. This supports the hypothesis that the poly-Q domain is instrumental in mediating pioneer activity, and that this activity contributes to embryo viability.

### The DNA-binding domain of GAF is necessary and sufficient for localization to foci

Contrary to our expectations, the poly-Q IDR was dispensable for GAF localization to subnuclear foci. We therefore sought to systematically identify the protein domains responsible for driving this distinctive localization. For this purpose, we generated transgenic fly lines that drove the expression of sfGFP-tagged GAF in the germline and early embryo and assayed for colocalization with endogenous mCherry-tagged GAF and mitotic retention during the MZT. We verified that transgenic expression of full-length GAF-sfGFP (1-519aa) recapitulated endogenous GAF localization both during mitosis and interphase (Figure 3A, S3A). As a control, we expressed sfGFP with the GAF nuclea localization signal (NLS) and confirmed that the protein was diffuse in the nucleus during interphase and was not mitotically retained (Figure 3A, S3B). Furthermore, transgenic expression of sfGFP-GAF without the poly-Q domain colocalized with endogenous GAF and was mitotically retained, as we also observed for endogenous sfGFP-GAF^SΔPQ^ (Figure 2C, 3A, S3A). Together, these controls support our use of transgenic expression to identify the domains of GAF required for nonuniform subnuclear distribution.

**Figure 3:**
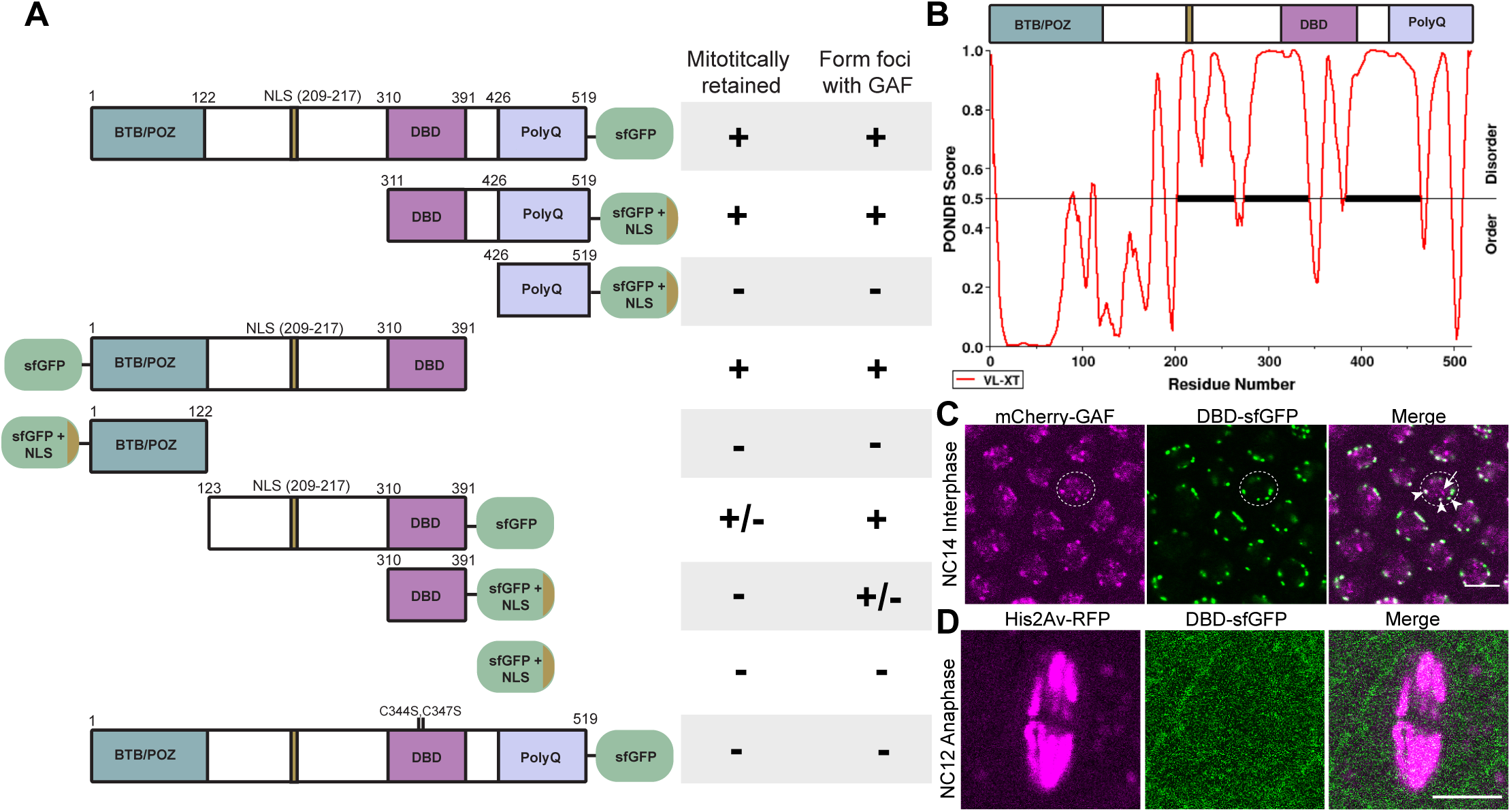
The DNA-binding domain of GAF is both necessary and sufficient for foci formation. A. Representations of tagged GAF truncations assayed (left) and whether or not those truncations were mitotically retained and localized to endogenous GAF foci (right). B. Prediction of intrinsically disordered regions of the short GAF isoform protein generated by PONDR. C. Interphase nuclei of an NC14 embryo expressing endogenously tagged mCherry-GAF and transgenically expressed DBD-sfGFP. mCherry-GAF is in magenta. DBD-sfGFP is in green. Scale bars, 5μM. A dotted circle indicates a representative nucleus. DBD-sfGFP colocalizes with mCherry-GAF (arrowheads), but there is a subset of mCherry-GAF foci that do not colocalize with DBD-sfGFP (arrow). D. Anaphase of a NC12-13 embryo expressing His2Av-RFP and the DBD-sfGFP transgene. His2Av-RFP is in magenta. DBD-sfGFP is in green. DBD-sfGFP is not retained on the mitotic chromosomes. Scale bars, 5μm.

Due to the reported role for IDRs in mediating protein aggregation, we used PONDR to identify other IDRs in the GAF protein outside of the poly-Q domain (Figure 3B). We identified an additional IDR N-terminal to the DNA-binding domain, suggesting that regions outside of the poly-Q domain might facilitate the localization of GAF to subnuclear foci and compensate for the absence of the poly-Q IDR. We therefore systematically made transgenes to express sfGFP tagged GAF lacking specific domains but retaining an NLS. All transgenes resulted in protein expression as assayed by imaging (Figure S3A, S3B). We discovered that all truncation proteins containing the intact DNA-binding domain (DBD) localized to foci (Figure 3A, S3A). Expression of the DBD alone localized to a subset of foci enriched for endogenous GAF (Figure 3A, C), demonstrating this domain alone is sufficient to drive localization to many GAF foci. Expression of proteins with either the poly-Q domain or the unstudied IDR from 123aa-310aa (IDR2) along with the DBD were able to localize to all foci marked by full-length GAF (Figure 3A, S3A). Thus, these IDRs may be important in mediating interactions for recruitment to a subset of GAF foci. In addition to being sufficient, the DBD domain was also necessary for localization. Full-length GAF containing point mutations in the single, DNA-binding zinc finger was not localized to foci, but instead was diffusely distributed in the nucleus (Figure 3A, S3B). Consequently, although the zinc finger mutant GAF was expressed in a background with endogenous wild-type GAF, the mutant protein was not recruited to GAF foci through protein-protein interactions. Our transgenic assays demonstrate that DNA-binding rather than protein-protein interactions is necessary for GAF to localize to subnuclear foci.

GAF is retained on mitotic chromosomes and is enriched at pericentric heterochromatin during mitosis (Raff et al. 1994). Using our transgenic system, we assayed for mitotic retention and demonstrated that despite localization to foci, the DBD alone was not mitotically retained (Figure 3A,C). However, the full-length zinc finger mutant GAF was also not retained during division (Figure 3A, S3B). Therefore, DNA-binding is necessary, but not sufficient for GAF localization to mitotic chromosomes. Addition of either the poly-Q domain or the entirety of the N-terminal portion of GAF to the DBD restored mitotic retention (Figure 3A, S3A). Expression of only the N-terminal IDR with the DBD resulted in a severe reduction in mitotic retention, despite the fact that this protein completely colocalized with endogenous GAF foci during interphase (Figure 3A, S3A). These data suggest that the N-terminal IDR and the poly-Q domain do not function equivalently to promote GAF binding during mitosis.

### GAF binds AAGAG satellite repeats in the early embryo

To identify the genomic regions that underlie the GAF foci, we leveraged the sfGFP-tagged DBD, which was sufficient for GAF localization to foci. We performed ChIP-seq using an anti-GFP antibody in stage 5 embryos expressing DBD-sfGFP. Despite successful immunoprecipitation of our spike-in material, there was not sufficient enrichment to call peaks from this dataset, including at regions bound by full-length GAF (Figure 4A). Based on the necessity of the DNA-binding domain for the localization of GAF to foci and our inability to detect enrichment in our DBD-sfGFP ChIP-seq dataset, we hypothesized that GAF foci might correspond to regions not represented in the reference genome, in particular the simple satellite AAGAG repeats enriched in *Drosophila* pericentric heterochromatin (Shatskikh et al. 2020). AAGAG repeats are abundant, comprising ∼6% of the *Drosophila* genome and would provide a highly concentrated region of the GA-repeat motif that GAF binds (Lohe and Brutlag 1987). Indeed, in third instar larval brain tissue GAF localizes to these repeats during mitosis but not during interphase (Platero et al. 1998). To identify whether GAF bound to these repetitive regions, we determined the percentage of total reads containing the (AAGAG)5 repeat in raw reads from the IP and input samples from our published GAF-sfGFP ChIP-seq data on stage 3 and stage 5 embryos (Gaskill et al. 2021). We then calculated the enrichment (IP/Input) of the percentage of reads that contained the AAGAG repeat. The AAGAG repeat was enriched in GAF IPs at both stages when compared to another pioneer factor that functions in the early embryo, Zelda (ZLD), which we would not expect to bind the AAGAG repeat (Gaskill et al. 2021)(Figure 4B). We next revisited our ChIP-seq data for the DBD alone to determine whether this domain was sufficient to bind to the repetitive AAGAG repeats despite the lack of detectable binding to euchromatic regions. This analysis identified enrichment of the AAGAG repeat in the DBD-sfGFP IPs at stage 5 at levels similar to full-length GAF (Figure 4B). We confirmed that the GAF foci localize to AAGAG repeats using DNA fluorescent in situ hybridization (FISH) on NC14 embryos using a (AAGAG)7 probe while simultaneously immunostaining for sfGFP-labelled GAF (Figure 4C). Together our data show that GAF binds to AAGAG repeats in the early embryo and at these regions the high local concentration of the GAF motif drives the formation of GAF foci.

**Figure 4.**
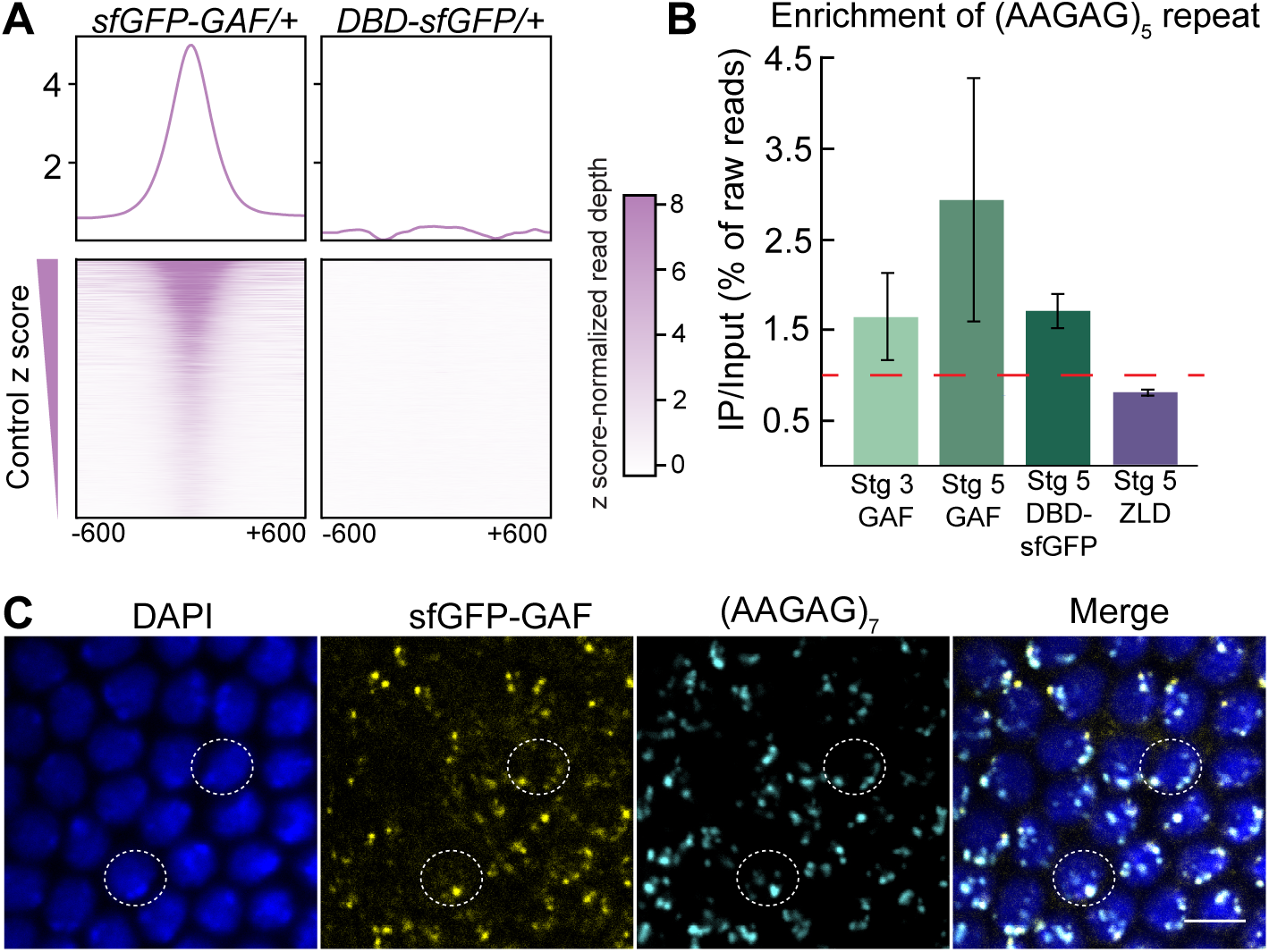
GAF foci correspond to the repetitive AAGAG element. A. Heatmaps of anti-GFP ChIP-seq performed on *sfGFP-GAF/+* embryos and embryos expressing transgenic DBD-sfGFP. B. The percentage of the total raw ChIP-seq reads containing (AAGAG)5 was determined and the ratio of the percentage of reads in the immunoprecipitation (IP) versus the input was plotted. Red line = IP/Input of 1. Error bars indicate the standard deviation of the two replicates tested. Stage 3 GAF-sfGFP, stage 5 GAF-sfGFP, and stage 5 Zld ChIP-seq datasets were analyzed from Gaskill et al. 2021. C. Images from DNA-FISH performed on NC14 *sfGFP-GAF(N)* homozygous embryos using an (AAGAG)7 probe. Embryos were immunostained with an anti-GFP antibody and labelled with DAPI. Dotted circles indicate representative nuclei. Scale bars, 5μm.

### A subset of GAF foci colocalize with HP1a

Satellite repeats are often silenced in the genome, however AAGAG repeats are expressed in various tissues throughout development, and AAGAG RNA is required for viability, the nuclear matrix, and sperm maturation (Mills et al. 2019; Pathak et al. 2013). Our data showed that GAF binds to AAGAG repeats in the early embryo, but the function of GAF at these repeats remained unclear. GAF has a variety of roles in transcriptional regulation, including activation, repression, insulator function, and chromatin organization (Adkins et al. 2006; Chetverina et al. 2021). To determine which of these many reported activities GAF may be employing at AAGAG repeats, we identified factors that colocalized with GAF at foci by focusing on proteins reported to be localized to foci. It has been well established that several types of foci form during interphase in the nucleus, including transcriptionally active hubs, insulator bodies, and heterochromatin domains (Rippe 2022; Maeda and Karch 2007; Peng et al. 2020). Since we had previously defined a role for GAF as a transcriptional activator during zygotic genome activation, we tested if GAF foci were sites of active transcription (Gaskill et al. 2021). Robust transcription initiates at the histone locus body (HLB) in two early detectable foci in the embryo (Chen et al. 2013; Huang et al. 2021; Hur et al. 2020). This phase-separated domain promotes high levels of histone gene expression and is marked by localization of the protein Multi Sex Combs (Mxc) (Hur et al. 2020). Using a GFP-tagged version of Mxc, we demonstrated that GAF is not localized to the HLB, consistent with previous data from fixed embryos (Figure S4) (Rieder et al. 2017). We then investigated if GAF foci were transcriptionally active hubs outside of the HLB. For this purpose, we imaged nascent transcription of a transgene driven by the regulatory region of the GAF-target gene *tailless* (*tll*) (Garcia et al. 2013). *tll* is zygotically expressed at NC14, bound by GAF at the promoter, and depends on GAF for transcription (Gaskill et al. 2021). We failed to observe strong colocalization between nascent transcription of this transgenic reporter and GAF foci (Figure S4). We propose that GAF foci are not transcription hubs, and that transcriptional activation is mediated by the population of GAF that is more diffuse in the nucleus.

In addition to its role in transcriptional activation, GAF functions as an insulator and interacts with several insulator proteins (Nègre et al. 2010; Lomaev et al. 2017; Wolle et al. 2015; Kyrchanova et al. 2018). Despite prior reports showing that GAF is in the Large Boundary Complex (LBC) with the insulator protein Mod (mdg4), we did not detect robust colocalization of these proteins (Figure S4). Nor did we identify colocalization with GAF and another insulator protein CTCF (Figure S4). GAF binds to Polycomb Response Elements (PREs) and interacts with subunits of the Polycomb repressive complex 2 (PRC2) (Ogiyama et al. 2018; Nègre et al. 2006; Strutt et al. 1997; Lomaev et al. 2017). As might be expected, we observed some colocalization of small GAF foci with Polycomb (Pc). However, the majority of GAF and Pc foci did not overlap (Figure S4).

Similar to other proteins classified as insulators, GAF is enriched at topologically associating domain (TAD) boundaries and Polycomb Response Elements (PREs), suggesting GAF might regulate 3D chromatin structure (Hug et al. 2017; Ogiyama et al. 2018). GAF has also been implicated in chromatin looping *in vitro* and *in vivo* (Batut et al. 2022; Mahmoudi et al. 2002; Petrascheck et al. 2005). To uncover changes in genomic contacts in the absence of GAF, we performed Hi-C on embryos in which GAF is degraded (GAF^deGradFP^) and control embryos at NC14 (Gaskill et al. 2021). The majority of 3D contacts were similar between the two conditions, indicating that GAF is not required for TAD formation during the MZT (Figure S5A). Chromatin is broadly divided into euchromatic (A) compartments, and heterochromatic (B) compartments (Maeshima et al. 2020), and we did not identify clear differences in this compartmentalization in the absence of GAF (Figure S5B). While the 3D organization of the genome remains largely unchanged in the absence of GAF, we did identify a small subset of loops that were lost when GAF was degraded, including at the *Antp* locus (Figure S5C). GAF binds to the anchors of this loop and has recently been implicated in tethering the enhancer to the promoter at this locus (Batut et al. 2022). Together our colocalization assays and Hi-C data show that GAF foci are unlikely to be either insulator or Polycomb bodies, and that GAF is not required for TAD formation or compartmentalization at NC14.

Our data suggest that GAF foci do not correspond to insulator bodies or transcription hubs. We therefore investigated whether GAF foci correspond to constitutive heterochromatin domains that are progressively formed during early embryogenesis. Heterochromatin protein 1 (HP1a) is thought to form phase separated domains in both *Drosophila* and mammals, and these domains contribute to the repression of heterochromatic regions through the selective concentration of silencing factors and exclusion of activating factors (Strom et al. 2017; Larson et al. 2017). We identified robust colocalization of sfGFP-GAF and HP1a tagged with RFP (Figure 5A). Despite this clear overlap at a subset of foci, in each nucleus there were GAF foci that did not contain HP1a, and HP1a foci that did not contain GAF (Figure 5A). Given the limited resolution of our confocal images, we used lattice light-sheet microscopy to determine the degree of colocalization more robustly between these two proteins. We confirmed the colocalization of GAF and HP1a, but the increased resolution clearly delineated subdomains within colocalized regions enriched for either GAF or HP1a (Supplemental movie S1). To determine whether GAF and HP1a colocalize at the AAGAG satellite repeats, we performed DNA FISH on NC14 embryos with a (AAGAG)7 probe and immunostained for HP1a. At NC14, both HP1a and the AAGAG repeats colocalize in the apical domains of the nuclei, reflecting the stereotypical Rabl configuration (Figure 5B). Some HP1a-enriched regions do not overlap the AAGAG repeat, likely representing additional heterochromatic regions. These data indicate that GAF forms foci by binding to AAGAG repeats and these correspond to heterochromatic genomic regions marked by HP1a.

**Figure 5:**
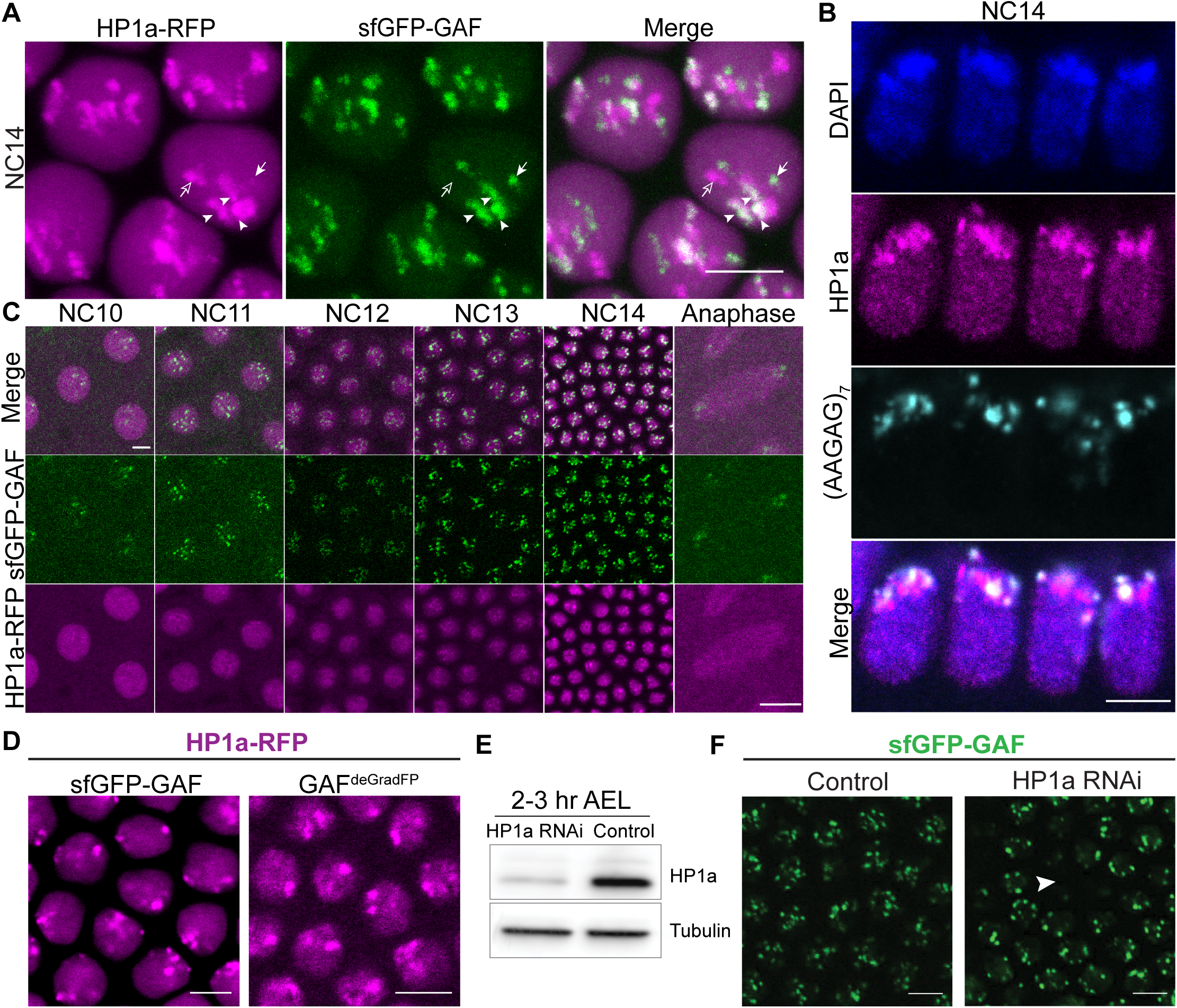
A subset of GAF foci localize with HP1a condensates. A. Representative image of interphase NC14 embryos laid by females expressing sfGFP-GAF and transgenic HP1a-RFP. HP1a-RFP is in magenta. sfGFP-GAF is in green. Scale bars, 5μM. Arrowheads indicate regions of colocalization. Closed arrow indicates sfGFP-GAF only foci. Open arrow indicates HP1a-RFP only foci. B. DNA-FISH performed on *sfGFP-GAF* embryos using an (AAGAG)7 probe. Embryos were immunostained with an anti-HP1a antibody and labelled with DAPI. Scale bars, 5μM. C. Images of a single embryo from cycles NC10-14 (indicated) laid by a female expressing endogenous sfGFP-GAF and transgenic HP1a-RFP. HP1a-RFP is in magenta. sfGFP-GAF is in green. Scale bars, 5μM. D. Representative images from interphase NC14 GAF^deGradFP^ and *sfGFP-GAF(N)* homozygous embryos laid by females expressing HP1a-RFP. HP1a-RFP is in magenta. Scale bars, 5μM. E. Immunoblot using anti-HP1a antibody on 2-3hr embryo extract from embryos expressing *Su(var)205* RNAi and controls. F. Representative images of NC14 embryos laid by females expressing endogenous sfGFP-GAF in a *Su(var)205* RNAi background. sfGFP-GAF is in green. Arrowhead highlights a region in which nuclei are below the plane of focus, indicative of the *Su(var)205* RNAi phenotype. Scale bars, 5μm.

Having established that many GAF foci correspond to HP1a-enriched AAGAG satellite repeats, we wanted to determine if GAF contributes to the formation of transcriptionally silent heterochromatin at these repeats. During the MZT, mature, constitutive heterochromatin domains are progressively established. It is not until NC13-14 that repetitive elements accumulate the repressive histone modification H3K9me3 and display the late replication characteristic of silenced regions (Yuan and O’Farrell 2016). HP1a, which binds to H3K9me3, begins to form small foci at NC11, and these HP1a domains do not mature into robust phase-separated domains until NC14 (Strom et al. 2017). In intriguing contrast, GAF foci can be clearly visualized by confocal microscopy starting at NC10. To determine the precise relationship between the timing of HP1a and GAF foci formation, we imaged sfGFP-GAF and HP1a-RFP in an embryo as it developed from NC10 through NC14. This analysis confirmed that GAF forms robust foci earlier than HP1a (Figure 5C, Supplemental movie S2). Because GAF is mitotically retained at pericentric heterochromatin while HP1a is not and forms foci prior to HP1a (Figure 5C, Supplemental movie S2), we hypothesized that GAF might be necessary for HP1a recruitment to foci. We tested if HP1a depended on GAF for foci formation by imaging HP1a-RFP in GAF^deGradFP^ embryos in which maternal GAF is degraded (Gaskill et al. 2021) (Figure 5D). In these embryos, HP1a foci were still clearly visible. We then tested if HP1a might be important for GAF foci formation by imaging sfGFP-GAF in embryos with HP1a knocked down by RNAi, and again did not detect any notable differences in foci formation (Figure 5E,F). Thus, while GAF and HP1a colocalize at AAGAG repeats, they can independently form subnuclear foci.

### GAF promotes H3K9me3 and represses transcription of AAGAG repeats

To determine the functional significance of GAF localization at AAGAG repeats, we tested if GAF regulated the establishment of H3K9me3, a histone modification instructive in the formation of HP1a-enriched constitutive heterochromatin. H3K9me3 is first detected at significant levels at NC14 (Yuan and O’Farrell 2016). Thus, GAF binding to AAGAG repeats at NC10 might be instrumental to the formation of heterochromatin through promoting H3K9me3. We first investigated the enrichment of H3K9me3 at GAF-bound AAGAG repeats during NC14 using DNA-FISH and immunostaining. We observed robust colocalization of many GAF foci with both AAGAG repeats and H3K9me3 signal at NC14 (Figure 6A). ChIP-seq for H3K9me3 at NC14 in control and GAF^deGradFP^ embryos showed a decrease of H3K9me3 at pericentric heterochromatin when GAF was degraded. (Figure 6B). Consistent with our Hi-C results, we observed that in the GAF^deGradFP^ embryos the Rabl conformation of the NC14 nucleus and the localization of AAGAG repeats to foci in the apical region was not disrupted, suggested GAF is not globally required for heterochromatin formation (Figure S5D). To more directly determine if the loss of H3K9me3 was specific to GAF-bound AAGAG repeats, we focused on the centromere of the 3^rd^ chromosome which is enriched for the dodeca satellite and has relatively few AAGAG repeats (Shatskikh et al. 2020). In this region, H3K9me3 signal was largely unchanged between control and GAF^deGradFP^ embryos, and loss of H3K9me3 is limited to sites that contain the AAGAG motif (Figure 6B). Because simple satellite repeats are not well represented in the reference genome assembly, we also analyzed the raw reads from the H3K9me3 ChIP-seq data. In GAF^deGradFP^ replicates there was a decrease in the enrichment of reads containing the AAGAG repeat compared to controls (Figure 6C). By contrast, when we performed the same analysis with the dodeca repeat, there was little difference between the GAF^deGradFP^ replicates and controls (Figure 6C). These data support a role for GAF in promoting the robust deposition of H3K9me3 at AAGAG repeats during the MZT.

**Figure 6:**
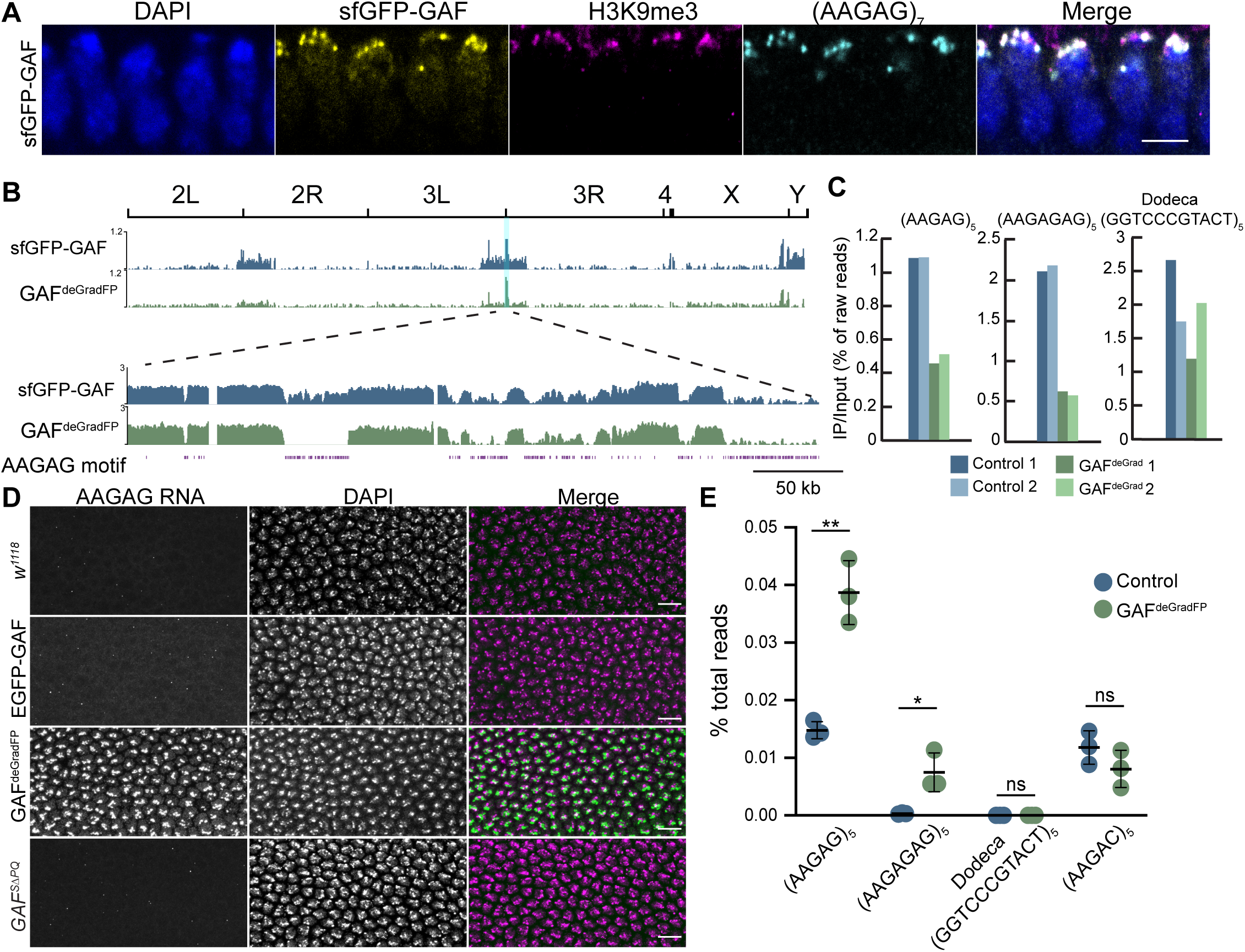
GAF is required to repress AAGAG satellite repeat expression during the MZT. A. DNA-FISH on *sfGFP-GAF(N)* homozygous embryos at NC14 using an (AAGAG)7 probe. Anti-GFP and anti-H3K9me3 antibodies were used for immunostaining. Scale bars 5μM. B. Genome browser tracks of IP read depth normalized to input from anti-H3K9me3 ChIP-seq performed on 2-2.5hr AEL *sfGFP-GAF(N)* homozygous and GAF^deGradFP^ embryos. The entire genome is shown. The region highlighted in blue from the 3^rd^ chromosome centromere is shown in detail below. C. IP/Input of the percentage of raw reads that contain the indicated satellite repeat sequences from anti-H3K9me3 ChIP-seq on control (*sfGFP-GAF(N)* homozygous) and GAF^deGradFP^ embryos at 2-2.5 hr AEL. D. RNA-FISH performed on *w^1118^*, *EGFP-GAF*, GAF^deGradFP^, and *GAF^SΔPQ^* embryos at NC14 using an (AAGAG)7 probe. Scale bars, 10μm. E. The percentage of unaligned total RNA-seq reads that contained the satellite repeat listed. Total RNA-seq was performed in 2-2.25 hr AEL *sfGFP-GAF(N)* homozygous and GAF^deGradFP^ embryos (Gaskill et al. 2021).

Our data suggest that GAF is concentrated at AAGAG satellite repeats and is instrumental in promoting heterochromatin formation. We therefore directly tested whether GAF was essential to silence transcription from AAGAG repeats. Failure to silence transcription from repetitive elements or establish heterochromatin at repetitive elements during the MZT leads to developmental defects (Ferree and Barbash 2009; Satyaki et al. 2014). To test the necessity of GAF for silencing repeat expression, we performed RNA FISH using an (AAGAG)7 repeat probe during NC14. We detected low levels of transcription from AAGAG repeats in control embryos, consistent with previous reports (Mills et al. 2019; Pathak et al. 2013) (Figure 6D). By contrast, in GAF^deGradFP^ embryos there was a robust increase in the RNA FISH signal from AAGAG repeats (Figure 6D). An increase in transcription of AAGAG repeats was not observed when we performed RNA FISH in embryos with only GAF^SΔPQ^ maternally deposited, indicating that GAF does not require the poly-Q domain to repress transcription from AAGAG repeats, separating pioneering activity from silencing of the repeats (Figure 6D). Analysis of total RNA-seq from GAF^deGradFP^ and control embryos at NC14 verified the increase in expression from AAGAG satellites in the absence of GAF (Gaskill et al. 2021). The percentage of total reads that contained (AAGAG)5 was significantly higher in GAF^deGradFP^ embryos compared to controls (Figure 6E). Additionally, the longer AAGAGAG repeat, which also contains a GAF motif, was significantly upregulated when GAF was degraded (Figure 6E). This effect was specific to GAF-bound repeats, as transcript abundance for several other simple repeats present in pericentromeric regions was unchanged in GAF^deGradFP^ embryos compared to controls (Figure 6E). The increased levels of AAGAG repeats observed are likely indicative of nascent transcription as RNA Pol II is robustly localized with the transcripts in GAF^deGradFP^ embryos (Figures S6). Together, our data demonstrate that GAF foci correspond to AAGAG satellites where GAF is concentrated by DNA binding to highly abundant GA-rich motifs and is required for robust methylation of H3K9 and transcriptional silencing.

## Discussion

To give rise to a new organism following fertilization, the chromatin of the specialized germ cells must be reprogrammed. During this reprogramming, both the active and silenced regions of the genome are established. We previously showed that in the embryo the pioneer factor GAF is required for activation of gene expression (Gaskill et al. 2021). Here, we demonstrated that in addition to activating the zygotic genome, GAF is required for silencing expression at satellite repeats where it is concentrated by high-density DNA motifs. Together these results provide insights into how a single factor has diverse roles in transcriptional regulation, likely through promoting the organization of chromatin into distinct subnuclear domains.

### GAF subnuclear domains are driven by DNA binding

Our data demonstrate that the formation of high-concentration GAF foci is driven by sequence-specific DNA binding rather than IDR-mediated multivalent interactions. Because GAF foci correspond to GA-rich satellite repeats, we propose that the high density of the GAF DNA-binding motif concentrates GAF at these genomic loci. While multivalent interactions mediated by IDRs drive protein aggregation in many systems (Boeynaems et al. 2018; Rippe 2022; Peng et al. 2020), recent work has begun to highlight the importance of additional domains to this process. For example, the DNA-binding domain of the human reprogramming factor KLF4 is both necessary and sufficient to form condensates in the presence of DNA (Sharma et al. 2021). DNA promotes surface condensation of KLF4, which can occur with a low saturation of molecules due to the local high density created at the DNA surface (Morin et al. 2022). Through similar but distinct mechanisms, nucleation at a specific DNA sequence is suggested to drive the formation of phase-separated domains, such as the histone locus body and nucleolus (Hur et al. 2020; Grob et al. 2014; Shevtsov and Dundr 2011). Our data support the importance of DNA binding in driving the formation of subnuclear regions of high protein concentration and highlights the complexity of mechanisms that can result in a nonuniform distribution of factors in the nucleus. This work reinforces that to determine if nonuniform protein distribution is driven by IDR-mediated phase separation it is essential to carefully analyze proteins expressed at endogenous levels and determine what portions of the protein are required for this distribution.

While we demonstrated that GAF DNA binding was required for localization to foci, we showed that not all endogenous GAF foci are occupied by the DBD alone. Either of the IDRs in combination with the DBD were enriched at all endogenous GAF foci, but the IDRs alone were unable to be recruited to foci. Indeed, even in the full-length GAF protein mutations in the DBD abrogated localization to foci. Recent single-molecule studies determined that regions outside the DBD, including the poly-Q IDR, are required for GAF to stably engage the genome (Tang et al. 2022). Thus, we propose that IDRs are essential for stabilizing GAF binding at a subset of loci, potentially through multivalent protein interactions.

### GAF supports heterochromatin establishment at AAGAG repeats

Despite the essential nature of heterochromatin, much remains enigmatic about how heterochromatin is established *de novo* during the MZT. The repressive histone modification H3K9me3 is not detectable until NC14, and HP1a does not form mature domains until the same time in development (Strom et al. 2017; Yuan and O’Farrell 2016). However, chromatin compaction is detected at satellite repeats beginning as early as NC8 (Shermoen et al. 2010). Furthermore, different satellite repeats accumulate H3K9me3 and HP1a at distinct time points in development and via different HP1a-recruitment mechanisms (Yuan and O’Farrell 2016; Seller et al. 2019). We demonstrated that GAF forms foci as early as NC10, prior to HP1a phase separation and H3K9me3 accumulation on chromatin. GAF foci are formed at AAGAG satellite repeats, and by NC14 these regions also colocalize with HP1a. This temporal relationship demonstrates that GAF binding to GA repeats precedes H3K9me3 deposition and HP1a enrichment. We propose that GAF binds GA-rich satellite repeats early in development and recruits heterochromatin factors to these regions. This is supported by the reduction in H3K9me3 and aberrant transcription of GA repeats in the absence of GAF. The recruitment of heterochromatin factors to GA-rich repeats is likely mediated through the BTB/POZ protein interaction domain of GAF, as we demonstrated that GAF continues to repress transcription from AAGAG repeats in the absence of the poly-Q domain.

GAF binds AAGAG repeats during interphase in the embryo, and mitotic retention of GAF to pericentric heterochromatin regions requires DNA-binding activity. Together these data suggest that GAF is bookmarking specific AAGAG binding sites throughout mitosis. The ability of GAF to remain bound to AAGAG targets throughout mitosis may contribute to the recruitment of heterochromatic factors to these regions during the rapid mitotic cycles that characterize early embryo development. A similar function was described for Prospero, a transcription factor that promotes neural differentiation through mitotic retention at pericentric heterochromatin, which facilitates efficient recruitment of heterochromatic factors such as HP1a upon mitotic exit (Liu et al. 2020). Together with prior data investigating the mechanisms of heterochromatin establishment, our data support a model whereby the timing of heterochromatin formation at distinct satellites may depend on sequence-specific binding factors that can recruit silencing machinery.

The role of GAF in heterochromatin formation is consistent with the nuclear defects observed in GAF-depleted embryos during the MZT (Gaskill et al. 2021). Anaphase bridges and micronuclei reminiscent of the division defects observed in GAF^deGradFP^ embryos are reported when D1 chromosomal protein (D1) and Proliferation disrupter (Prod) are depleted (Jagannathan et al. 2019, 2018). D1 and Prod bind sequence specifically to repeats and function to spatially condense and organize repetitive regions in the genome, allowing for the formation of the chromocenter. Based on this evidence, it has been suggested that although there is little sequence conservation between closely related species, satellite repeats have a conserved function in recruiting proteins that facilitate the bundling of heterochromatin from multiple chromosomes into organized subnuclear domains (Jagannathan et al. 2019, 2018). AAGAG repeats continue to localize in foci at the heterochromatic apical regions of nuclei in NC14 GAF-depleted embryos, and Hi-C data indicates that the loss of GAF does not affect organization of heterochromatin and euchromatin into distinct subnuclear compartments. This suggests that GAF is not essential for global genome organization. GAF might have a role at repeats separate from genome organization, or another factor, such as D1 and Prod, may function redundantly with GAF to preserve spatial organization of repetitive regions when GAF is depleted. This redundancy is supported by the fact that D1 and Prod are partially redundant to each other in chromocenter formation (Jagannathan et al. 2019). Alternatively, GAF knockdown may compromise spatial organization of repetitve regions in a subset of embryos, and this may cause catastrophic failure of nuclear divison and embryo death prior to NC14. In our analysis, we selected for NC14 embryos, as this is when distinctive marks of heterochromatin are clear. This resulted in the exclusion of dying embryos with dramatic defects in nuclear division, which might be the result of perturbed chromatin organization due to the absence of GAF. Together our data support an essential role for GAF in the formation of heterochromatin at AAGAG satellite repeats, but additional studies will be required to determine the role of GAF in the maintenance of a prolonged silenced state. These future studies will have important implications for our understanding of heterochromatin formation more generally as the role of GAF in heterochromatin formation may be conserved. Notably, cKrox, the mouse ortholog of GAF, drives localization of GA-rich loci to heterochromatic domains on the nuclear periphery (Zullo et al. 2012).

### Mechanisms of distinct GAF functions

Here, we identified a novel role for GAF in repression of satellite repeats. This was unexpected as our prior work, and that of others, had largely focused on the role of GAF as a pioneer factor essential for activating transcription (Gaskill et al. 2021; Fuda et al. 2015; Judd et al. 2021; Tang et al. 2022; Bellec et al. 2018; Tsukiyama et al. 1994). Together our studies demonstrate that in a single nucleus at one point in development, GAF is functioning to establish both the silent and active transcriptional state. How GAF can perform these two opposing functions remains unclear but based on our data we propose that regions of high GAF density result in transcriptional silencing. By contrast, more diffuse populations of GAF may promote transcriptional activation. This model of protein density driving changes in transcription-factor function is exemplified by the oncogenic transcription factor EWS::FLI1. EWS::FLI1 can function as either an activator or a repressor depending on the level of low-complexity domain interaction (Chong et al. 2022). At endogenous levels, EWS::FLI1 forms hubs at activated target genes. Induction of phase separation at EWS::FLI1 hubs by overexpression of low-complexity domains results in the silencing of genes normally activated by EWS::FLI1. The authors suggested a model where hub formation is precisely tuned for target-gene activation, with too few or too many IDR-mediated multivalent interactions resulting in gene repression. Therefore, the high concentration of GAF at satellite repeats may similarly result in repression, while lower concentrations of GAF promote transcription.

GAF interacts with a variety of co-factors that function in transcriptional activation and repression (Lomaev et al. 2017). It is possible increased protein concentration results in repression via the selective trapping or exclusion of specific cofactors. For example, the high density of GAF at GA-repeats may result in the sequestration of repressive factors in these foci and the exclusion of activating factors. Indeed, phase separated domains, like heterochromatic HP1a, have been demonstrated to have this functionality (Strom et al. 2017). GAF is also post-translationally modified, and this could provide another level of regulation to control GAF function. GAF is phosphorylated, O-glycosylated, and acetylated (Bonet et al. 2005; Jackson and Tjian 1988; Aran-Guiu et al. 2010). These modifications might selectively regulate the ability of GAF to interact with co-factors or directly influence recruitment of GAF to specific subnuclear domains. Human HP1*α* requires phosphorylation for phase separation, demonstrating the importance of post-translational modifications in regulating protein localization (Larson et al. 2017).

Our data revealed a role for GAF in establishing heterochromatin in the early embryo. We demonstrate that GAF is concentrated at satellite repeats that contain a high density of the GAF DNA-binding motif and that GAF binding at these sites results in silencing of gene expression. Along with prior studies, we show that GAF is distinctive as it functions broadly in both transcriptional activation and repression during the dynamic genomic reprogramming required for early development. We propose that the non-uniform distribution of GAF in the nucleus enables it to have opposing transcriptional roles within a single nucleus. Together, our data support a model in which a subset of transcription factors are important for organizing the nucleus into functionally distinct domains based on their subnuclear concentration, and this is essential for regulating gene expression during dramatic changes in cell fate.

## Materials and Methods

### *Drosophila* strains and genetics

All stocks were grown on molasses food at 25°C. Fly strains used in this study: *w^1118^*, *w*;*His2Av-mRFP (III)* (Bloomington *Drosophila* Stock Center (BDSC) #23650), *w*;*mat-α-GAL4-VP16 (II)* (BDSC #7062), *w*;*Trl^s2325^/TM3* (BDSC #12088*), yw; P{CTCF-GFP.FPTB}* (BDSC #64810), *w;PBac{GFP-mod(mdg4).S}* (BDSC #51351), *yw;PBac{mxc-GFP.FPTB}* (BDSC # 84130), *w; P{Pc-EGFP}* (BDSC #9593), *His2AV-RFP(II);MCP-GFP (III)* (Garcia et al. 2013), *UAS-dsRNA-Su(var)205* (BDSC #36792), *HP1a-RFP* (II) (Lipsick, G. Karpen), *sfGFP-GAF (N)* and *nos-degradFP (II)* were generated previously by our lab and are described in Gaskill et al. 2021.

#### Transgenic lines

GAF truncation GFP-tagged transgenic flies were made using the sequence from the short GAF isoform (519aa). The NLS of GAF (209-217aa) was added to the C-terminus of sfGFP for the truncations that lacked the endogenous NLS. Full length (1-519aa)-sfGFP, DBD-PQ (311-519aa)-sfGFP, PQ (426-519aa)-sfGFP, sfGFP-BTB/POZ-DBD (1-391aa), sfGFP-BTB/POZ (1-122aa), IDR-DBD (123-391aa)-sfGFP, Full-length zinc finger mutant (1-519aa, C344S, C347S)-sfGFP were made by PhiC31 integrase-mediated transgenesis into the PBac{yellow[+]-attP-3B}VK00037 docking site (BDSC #9752) by BestGene Inc. DBD (310-391aa)-sfGFP was made by PhiC31 integrase-mediated transgenesis into attP40 (25C6) docking site by BestGene Inc. All transgenes were cloned using Gibson assembly into an attB vector with the *nanos* promoter and 5’UTR that was used to generate transgenes in Gaskill et al. 2021.

To generate the MS2 transgene driven by the *tll* promoter, 3.3kb of the *tll* regulatory region was cloned upstream of 24x MS2 loops using Gibson assembly into an attB vector. Transgenes were made by PhiC-mediated Recombinase Mediated Cassette Exchange (RMCE) into the P{attP.w[+].attP}JB38F docking site (BDSC #27388) by BestGene Inc.

The following GAF mutant alleles were generated using Cas9-mediated genome engineering (outlined in detail below): GAF^L^, GAF^SΔPQ^, mCherry-GAF, and sfGFP-GAF^SΔPQ^

To obtain the embryos for microscopy in a *Su(var)205* knockdown background, we crossed *mat-α-GAL4-VP16 (II)/CyO, sfGFP-GAF (N)(III)* to *UAS-dsRNA-Su(var)205* (BDSC #36792) (II). Resulting *mat-α-GAL4-VP16/UAS-dsRNA-Su(var)205 (II), sfGFP-GAF (N)/+(III)* females were crossed to their siblings, and their embryos were collected.

### Cas9-genome engineering

Cas9-mediated genome engineering as previously described (Hamm et al. 2017) was used to generate the N-terminal mCherry-tagged GAF. The double-stranded DNA (dsDNA) donor was created using Gibson assembly (New England BioLabs, Ipswich, MA) with 1 kb homology arms flanking the mCherry tag and GAF N-terminal open reading frame. The mCherry sequence was placed downstream of the GAF start codon. A 3xP3-DsRed cassette flanked by the long-terminal repeats of PiggyBac transposase was placed in the second GAF intron for selection. The guide RNA sequence (TAAACATTAAATCGTCGTGT) was cloned into pBSK with U63 promoter using inverse PCR. Purified plasmid was injected into embryos of *yw; attP40{nos-Cas9}/CyO* by BestGene Inc. Lines were screened for DsRed expression to verify integration. The entire 3xP3-DsRed cassette was cleanly removed using piggyBac transposase, followed by sequence confirmation of precise tag integration.

To generate *GAF^L^*, *GAF^SΔPQ^*, *sfGFP-GAF^SΔPQ^* lines, single-stranded oligodeoxynucleotide (ssODN) donors containing the desired mutations were produced by Integrated DNA Technologies. To generate *GAF^L^*, a stop codon was introduced at the beginning of the 6^th^ exon of the long isoform. Immediately downstream of the stop codon a HindIII site (AAGCTT) was generated, and additional mutations were made in the seed region of the guide site in the 5^th^ GAF intron to prevent Cas9 nuclease recutting. To generate *GAF^SΔPQ^*, a 297bp deletion of exon 5 was created, removing sequence unique to the short isoform. To generate the deletion, two guide sites were used flanking the 297bp deletion. The deletion disrupted one guide site, and a mutation was created in the seed region of the other guide site in the 3’UTR of the short isoform. The guide RNA sequences (GAF^L^ - GCGGCAGTCTTCTCACCAGC) (GAF^SΔPQ^ - AGCCTTCAATCATTCCAACG and ACGAGAGTGATATCGAATGC) were cloned into pBSK with the U63 promoter using inverse PCR. The ssODN and guide RNA plasmids were injected into embryos of *yw; attP40{nos-Cas9}/CyO* by BestGene Inc. Lines were screened using PCR and HindIII digestion for GAF^L^, and PCR screening for the 297bp deletion for GAF^SΔPQ^. The regions were then sequenced to confirm mutation without errors. *sfGFP-GAF^SΔPQ^* was created identically to *GAF^SΔPQ^* except the guide plasmids and ssODNs were injected into *nos-Cas9 (II)/sfGFP-GAF(N) (III)* embryos provided to BestGene Inc.

In both *sfGFP-GAF^SΔPQ^* and *GAF^SΔPQ^* lines, the 297bp deletion included the short isoform stop codon. Both mutant lines create a truncated protein product. Based on our sequencing of the mutant lines, we have predicted the additional amino acid sequence that would be translated from the transcript until the closet stop codon.

GAF^SΔPQ^: 424 – PPPAEPSIIPTHQRHHHPHFQKNIKKKNITLTKTICK* sfGFP-GAF^SΔPQ^: 423 – THLQPSLQSFQRTNDTIIHISKKTLKKKT*

### Confocal microscopy

Embryos were dechorionated in 50% bleach for 2 min and subsequently mounted on a hydrophobic membrane coated in heptane glue. Embryos were covered in halocarbon 27 oil prior to the addition of a coverslip. Embryos were imaged on a Nikon A1R+ confocal using a 100x objective at the University of Wisconsin-Madison Biochemistry Department Optical Core. Nuclear density, based on the number of nuclei/2500 μm^2^, was used to determine the cycle of pre-gastrulation embryos. Nuclei were marked with His2AV-RFP. Image J (Schindelin et al. 2012) was used for post-acquisition image processing. For all images a single z-plane is shown, except Figure 5A, which is a maximum intensity projection of multiple z-stacks.

### Lattice light-sheet microscopy data acquisition and analysis

Lattice light-sheet microscopy was performed as described previously (Mir et al. 2018a). A 5mm glass coverslip was rendered adhesive by deposition of a small drop of glue solution prepared by dissolving a roll of double-side scotch tape in heptane. The glue solution was allowed to completely dry before embryos were introduced. Embryos were collected from cages over a 90 minute laying period and arranged on a 5mm diameter glass cover slip. Lattice Light-Sheet Microscopy was performed using a home-built implementation of the instrument following designs from the Betzig Lab (Chen et al. 2014). A 30 beam square lattice was generated with inner and outer numerical apertures of 0.505 and 0.60 respectively. The sheet was dithered over a 5 µm range in 200 nm steps during each exposure to create a uniform excitation profile. 488 nm and 561 nm lasers were used to excite GFP and RFP respectively, with laser powers measured to be 900 µW for 488 nm and 560 µW for 561 nm at the back aperture of the detection objective. Two Hamamatsu ORCA-Flash 4.0 digital CMOS cameras (C13440-20CU) were used for detection. An image splitting long-pass dichroic (Semrock FF-560) was placed in between the two cameras to separate emission wavelengths of over and under 560 nm, bandpass filters Semrock FF01-525/50 for GFP and Semrock FF01-593/46 for RFP were placed in front of the cameras. Images at each excitation wavelength were acquired sequentially at each z-plane with an exposure time of 100 ms for each channel and a 5 second pause between volume acquisitions, these settings resulting in 21 second interval between volumes. Data were rendered using Imaris with no further processing for visualization and quantified as described below.

The HP1-RFP channel was used to segment nuclei in 3D. The nuclei images were convolved with a difference of Gaussians filter, followed by a threshold calculated using Otsu’s method to generate a binary mask. The mask was cleaned to remove noise and any nuclei touching the image border. Nuclei were then linked across frames using a 3D nearest-neighbor algorithm. Nuclei masks and trajectories were manually inspected for accuracy. Only nuclei tracked for at least 90% of each nuclear cycle were retained for further analysis.

To identify the volume occupied by GAF foci, a 93-percentile threshold calculated across all time points was applied to the GAF-GFP channel within the nuclei. Volumes were calculated from the volume of each image voxel (0.1x0.1x0.2 um^3) within the binary foci and nuclear masks.

To count discrete GAF puncta, the GAF-GFP channel was convolved with a Gaussian filter (sigma=1.2) and local maxima within a 5-voxel neighborhood were identified. These puncta were then filtered using the GAF foci volumetric mask. Images were manually inspected to ensure that this filter identified all visually apparent foci within each nucleus. Analysis code is available here: https://gitlab.com/mir-lab/publications/gaf-llsm-analysis

### Chromatin immunoprecipitation

Chromatin immunoprecipitations were performed as described previously (Blythe and Wieschaus 2015). ChIP was performed using the anti-GFP antibody (abcam 290) on embryos from *sfGFP-GAF^SΔPQ^* heterozygous females and *nos-DBD-sfGFP* heterozygous females. ChIP was performed using the anti-H3K9me3 antibody (Active motif 39161) on *sfGFP-GAF(N)* homozygous and GAF^deGradFP^ embryos.

Briefly, 400 embryos from 2-2.5 hr lays were collected, dechorionated in 50% bleach for 3 min, fixed for 15 min in 4% formaldehyde and hand-sorted by morphology to ensure they were stage 5. Embryos were then lysed in 1 mL of RIPA buffer (50 mM Tris-HCl pH 8.0, 0.1% SDS, 1% Triton X-100, 0.5% sodium deoxycholate, and 150 mM NaCl). The fixed chromatin was sonicated for 20 s 11 times at 20% output and full duty cycle (Branson Sonifier 250). At this point sheared spike-in chromatin from H3.3-GFP mouse cells (sfGFP-GAF^SΔPQ^ and DBD-sfGFP ChIP) or *D. virilis* embryos (H3K9me3 ChIP) was added to the sonicated chromatin. Chromatin was incubated with 6 μg of anti-GFP antibody (abcam #ab290) or 10 μl anti-H3K9me3 antibody (Active Motif, # 39162) overnight at 4°C and then bound to 50 μl of Protein A magnetic beads (Invitrogen). The purified chromatin was washed, eluted, and treated with 90 μg of RNaseA (37°C, for 30 min) and 100 μg of Proteinase K (65°C, overnight). The DNA was purified using phenol/chloroform extraction and concentrated by ethanol precipitation. Each sample was resuspended in 25 μl of water. Sequencing libraries were made using the NEB Next Ultra II library kit. For sfGFP-GAF^SΔPQ^ ChIP-seq, libraries were sequenced on the Illumina NextSeq 500 using 75bp single-end reads at the Northwestern Sequencing Core (NUCore). For DBD-sfGFP ChIP-seq, libraries were sequenced on the Illumina HiSeq 4000 using 50bp single-end reads at the Northwestern Sequencing Core (NUCore). For H3K9me3 ChIP-seq, libraries were sequenced on the Illumina NovaSeq 600 using 150bp paired-end reads at the UW Madison Biotechnology Center.

### ChIP-seq Data Analysis

Pearson’s correlation coefficients were calculated for all ChIP-seq replicates using deeptools (Ramírez et al. 2016) and are reported in Table S3.

#### sfGFP-GAF^SΔPQ^ and DBD-sfGFP ChIP

ChIP-seq data was aligned to a combined *Drosophila melanogaster* reference genome (version dm6) using bowtie 2 v2.3.5 (Langmead and Salzberg 2012) with the following non-default parameters: -k 2, --very-sensitive. Aligned reads with a mapping quality < 30 were discarded, as were reads aligning to scaffolds or the mitochondrial genome. To identify regions that were enriched in immunoprecipitated samples relative to input controls, peak calling was performed using MACS v2 (Zhang et al. 2008) with the following parameters: -g 1.2e8, --call-summits. To focus analysis on robust, high-quality peaks, we used 100 bp up- and downstream of peak summits, and retained only peaks that were detected in both replicates and overlapped by at least 100 bp. All downstream analysis focused on these high-quality peaks.

To compare GAF-binding sites at in *sfGFP-GAF^SΔPQ^* to *sfGFP-GAF* controls, we used intersectBed from Bedtools (Quinlan and Hall 2010) to compare different sets of peaks. Peaks overlapping by at least a 20bp overlap were considered to be shared. DeepTools (Ramírez et al. 2016) was used to generate read depth for 10 bp bins across the genome. A z-score was calculated for each 10 bp bin using the mean and standard deviation of read depth across all 10 bp bins. Z-score normalized read depth was used to generate heatmaps, metaplots, and genome browser tracks.

#### H3K9me3 ChIP

ChIP-seq data was aligned to the *Drosophila melanogaster* reference genome (version dm6) using bowtie 2 v2.3.5 (Langmead and Salzberg 2012) without the following default parameters to retain multimapping reads: --no-mixed --no-discordant. All aligned reads were kept regardless of mapping quality to retain multimapping repeats. bamCompare was used with the following parameters --scaleFactorsMethod SES, --operation log2, to generate bigWig files of the H3K9me3 signal normalized to the input.

#### AAGAG repeat ChIP analysis

To analyze raw reads for simple satellite repeats raw fastq files were converted to fasta files. Fasta files were then searched for (AAGAG)5 and the reverse complement sequence. To determine how well the repeat of interest was immunoprecipitated, we calculated the percentage of total reads that contained the repeat of interest in both the IP and paired input raw reads. The IP/input was then calculated. A scaling factor calculated from the spike-in normalization was applied to the IP/input for the H3K9me3 ChIP-seq replicates. To verify that the *D. virilis* or mouse spike in chromatin would not confound our analysis, we determined the amount of reads with (AAGAG)5 in input samples from *D. virilis* ovaries (Le Thomas et al. 2014) and MEF cells (Rodier et al. 2015). In contrast to the *D.melanogaster* ChIP-seq input samples analyzed for this study in which 0.39 - 3.28% of total reads contained the (AAGAG)5 repeat, in *D.virilis* input material only 0.00059 - 0.00064% of the total reads contained the (AAGAG)5 repeat, indicating this repeat is not abundant in the *D.virilis* genome and would not impact our analysis. Similarly, in MEF input samples only 0.0032 – 0.0042% of total reads contain the (AAGAG)5 repeat.

### ChIP-seq spike-in normalization

#### sfGFP-GAF^SΔPQ^ and DBD-sfGFP ChIP

Prior to addition of the anti-GFP antibody, mouse chromatin prepared from cells expression an H3.3-GFP fusion protein was added to *Drosophila* chromatin at a 1:750 ratio. Following sequencing, reads were aligned to a combined reference genome containing both the *Drosophila* genome (version dm6) and the mouse genome (version mm39). Only reads that could be unambiguously aligned to one of the two reference genomes were retained. To control for any variability in the proportion of mouse chromatin in the input samples, the ratio of percentage of spike in reads in the IP relative to the input were used. A scaling factor was calculated by dividing one by this ratio. Z-score normalized read depth was adjusted by this scaling factor, and the resulting spike-in normalized values were used for heatmaps.

#### H3K9me3 ChIP

Prior to addition of the anti-H3K9me3 antibody, *D. virilis* chromatin prepared from stage 5 embryos was added to the *D. melanogaster* chromatin at a 1:25 ratio. Following sequencing, reads were aligned to a combined reference genome containing both the *D. melanogaster* (version dm6) *D. virilis* genome. Reads were aligned to the combined reference genomes using parameters that retained multimapping reads. To control for any variability in the proportion of *D. virilis* chromatin in the input samples, the ratio of percentage of spike in reads in the IP relative to the input were used. A scaling factor was calculated by dividing one by this ratio. The scaling factor was used to adjust the IP/input value in the analysis of the raw reads.

### Total RNA-seq analysis

To analyze raw reads for simple satellite repeats raw fastq files were converted to fasta files. Fasta files were searched for x5 repeats of common simple satellites and their reverse complement sequences. To determine if the repeat of interest changed in expression between GAF^deGradFP^ and control embryos, we calculated the percentage of total reads that contained the repeat of interest in each replicate. A two-tailed T-test was used to determine significance between the percentage of repeat reads in the GAF^deGradFP^ compared to control replicates.

### Adult phenotyping and viability assays

*GAF^SΔPQ^* and *GAF^L^* mutant alleles were assayed in *trans* to the GAF null mutation *Trl^s2325^* to verify that any identified phenotypes were the result of a mutation in *Trl* and not a background mutation on the same chromosome. Trans-heterozygous adult progeny were checked for phenotypes and crossed to *w^1118^* to determine fertility.

For the viability assays, three to five heterozygous males and five to 10 heterozygous females of the indicated genotypes were mated in standard molasses vials with dry yeast and flipped twice at 2-day intervals. Two days after the final flip, the adult flies were cleared from the vials and their progeny were allowed to reach adulthood. The *Trl^s2325^* allele was used as the GAF null. Over 900 adults were counted for each cross. The ratio of TM3 and non-TM3 adults was determined and the *χ*2 value was calculated for each cross, correcting for the observed ratio from the GAF null/wild-type cross.

### Hatching-rate assays

A minimum of 50 females and 25 males of the indicated genotypes were allowed to mate for at least 24 hours before lays were taken for hatching rate assays. Embryos were picked from overnight lays and approximately 200 were lined up on a fresh plate. Unhatched embryos were counted 26 hours or more after embryos were picked.

### Antibody generation and purification

An N-terminal GAF antibody recognizing the first 130 amino acids of GAF was used for immunoblotting. To generate these antibodies, rabbits were immunized by Covance, Inc., with maltose binding protein (MBP) fused to amino acids 1–130 of GAF and purified against the same portion of the protein fused to glutathione S-transferase (GST). Similar to other anti-GAF antibodies (Benyajati et al. 1997; Bhat et al. 1996), this antibody recognizes the short GAF isoform band at approximately 70 kD in an immunoblot on *w^1118^* overnight embryo extract (Figure S2E).

### Whole-embryo immunostaining

Embryos were dechorionated and added to 4% formaldehyde in 1x PBST (0.1% Triton-X) with an equal volume of heptanes. Embryos were rocked for 20 minutes at room temperature. The aqueous layer was removed, and methanol was added. Embryos were vortexed for 30 seconds, and all liquid was removed after the embryos settled. Embryos were washed 3x with methanol and 3x with 100% ethanol and stored in ethanol at -20C until use. To rehydrate embryos, they were washed sequentially for 10 min at room temp in 50% EtOH 50% PBST, 25% EtOH 75% PBST, and 100% PBST. They were then incubated overnight at 4C with the primary antibodies in PBST (Active motif (2AG-6F12-H4) anti-H3K9me3 (1:100)) (Abcam 290 anti-GFP (1:500)). The next day embryos were washed 3x for 5 minutes in PBST and incubated for 1.5 hours at room temp with secondary antibodies (Dylight 550 goat anti mouse 1:400 and Dylight 488 goat anti rabbit 1:400). Embryos were washed 3x for 5 minutes in PBST, then for 10 minutes in PBST + DAPI. Embryos were finally washed in for 1 minute in PBS and mounted in 70% glycerol and 1x PBS.

### Immunoblotting

Proteins were transferred to 0.45 μm Immobilon-P PVDF membrane (Millipore) in transfer buffer (25 mM Tris, 200 mM Glycine, 20% methanol) for 60 min at 500mA at 4°C. The membranes were blocked with blotto (2.5% non-fat dry milk, 0.5% BSA, 0.5% NP-40, in TBST) for 30 min at room temperature and then incubated with anti-GAF (1:250, this study), anti-GFP (1:2000, Abcam #ab290, #ab6556), anti-Tubulin (DM1A) (1:5000 Sigma #T6199), anti-HP1a (1:50, DSHB, C1A9) overnight at 4°C. The secondary incubation was performed with goat anti-rabbit IgG-HRP conjugate (1:3000 dilution, Bio-Rad #1706515) or anti-mouse IgG-HRP conjugate (1:3000 dilution, Bio-Rad # 1706516) for 1 hour at room temperature. Blots were treated with SuperSignal West Pico PLUS chemiluminescent substrate (Thermo-Scientific) and visualized using the Azure Biosystems c600 or Kodak/Carestream BioMax Film (VWR).

### cDNA screening

Ten overnight embryos from the indicated genotype were picked into Trizol (Invitrogen

#15596026) with 200 μg/ml glycogen (Invitrogen #10814010). RNA was extracted and cDNA was generated using Superscript IV (Invitrogen). cDNA was diluted 1:10 and used for PCR. Two primer sets were used. Primer set 1 (F primer-CCTTTCTGCTGGACTTGCTAAAG, R primer-CGGATTGTGCCACCAGTT) amplified both the long and short isoform transcripts. With these primers, the long isoform transcript band is 1522 bp, the WT short isoform band is 2034 bp, and the truncated short isoform band is 1737 bp. Primer set 2 (F primer-CGACCAAGACCAACTGATTGC, R primer-GAACACAAATCATTCGATCAGATC) amplified only the short isoform transcript. With these primers the WT short isoform band is 1310 bp and the truncated short isoform band is 1013 bp. Bands marked with an asterisk were excised, purified, and sequenced to confirm they were the short or long transcripts.

### Hi-C experimental procedure and analysis

500 *sfGFP-GAF(N)* homozygous or GAF^deGradFP^ hand sorted 2-2.5 hr AEL embryos were used as input for each replicate. Hi-C experiments and initial data processing were performed as described previously (Stadler et al. 2017) with minor modifications: the restriction enzyme MluCI (^AATT, NEB R0538L) was used, as we found that it gives more even coverage of the AT-rich Drosophila genome than GATC-cutters, and IDT xGen adaptors were used in place of Illumina adaptors. Insulation scores were computed by first computing the directionality (ratio of contacts with bins to the right vs. left) for each 500 bp bin in the genome, then performing a rolling difference calculation with window and step sizes of 16 bins (8 kb), followed by smoothing with a moving average with a 5 kb window size. Compartment scores were calculated for 25 kb bins of whole chromosomes by normalizing each bin for distance from the diagonal (i.e., observed/expected), calculating the covariance matrix, and taking the first eigenvector. Python scripts for all analyses are available at https://github.com/michaelrstadler/hic

### DNA fluorescent in-situ hybridization

#### Fixation

DNA FISH was performed on 1.5-3 hr AEL embryos as described previously (Bantignies and Cavalli 2014). Briefly, embryos were dechorionated and transferred to buffer A (60mM KCl, 15 mM NaCl, 0.5 mM spermidine, 0.15 mM spermine, 2 mM EDTA, 0.5 mM EGTA and, 15 mM PIPES at pH7.4 made fresh) with 4% paraformaldehyde. Equal amount of heptane was added followed by 25 minutes of incubation on an orbital shaker at max speed. Fixed embryos were devitellinized, washed twice in 100% methanol, and stored at -20°C.

#### Hybridization

Embryos were rehydrated and incubated with 200 µg/ml RNAse A in PBT at 4°C on a rotating wheel overnight. Next day, embryos were incubated in PBS with 0.3% Triton X-100 (PBS-Tr) for 1 hour before being gradually acclimated to 100% pre-hybridization mixture (pHM) (50% Formamide, 4x SSC, 100 mM NaH2PO4, pH7.0, and 0.1% Tween 20). Once in 100% pHM, embryos were incubated for 15 min at 80°C. The DNA probe 5Cy5/(AAGAG)7 was denatured in FISH hybridization buffer (FHB) (50% Formamide, 10% Dextran sulfate, 2x SSC, Salmon sperm DNA 0.5 mg/ml (0.05%)) for 10 minutes at 90°C. Samples hybridized to the probe overnight in a thermomixer set to 37°C with 450 rpm of agitation. Embryos were washed in (1) 50% Formamide, 2x SSC, and 0.3% CHAPS x 2; (2) 40% Formamide, 2x SSC, and 0.3% CHAPS; (3) 30% Formamide, 70% PBT; and (4) 20% Formamide, 80% PBT for 20 minutes each in a thermomixer set to 37°C and 850 rpm. Washes continued with 10% Formamide, 90% PBT, 100% PBT, and 100% PBS-Tr for 20 minutes each at room temperature on a rotating wheel.

#### Immunostraining

Embryos were processed for immunostaining by first incubating on a rotator in blocking solution of 3% BSA in PBS-Tr for two hours at room temperature. Primary antibodies were diluted in blocking solution (1:500 Rabbit anti-GFP Abcam 290; 1:50 Mouse anti-HP1a DSHB, C1A9) and incubated with embryos at 4°C overnight. Embryos were then washed in PBS-Tr 3x for 5 minutes and 3x for 20 minutes at room temperature while rotating in between each wash. Secondary antibody was diluted at 1:1000 in blocking solution (Goat Anti-Rabbit IgG DyLight 488 conjugated; Goat Anti-Mouse IgG Dylight 550 conjugated) and allowed to incubate with the embryos for 1 hour at room temperature. Subsequent washing in PBS-Tr was the same as for the primary antibodies. DAPI (4’,6-diamidino-2-phenylindole; Invitrogen REF: D1306) diluted at 1:1000 in PBT was added to the samples for 10 minutes at room temperature on a rotating wheel followed by a wash in PBT for 10 minutes. Embryos were mounted in 70% glycerol in PBS.

### RNA fluorescent in-situ hybridization

RNA *in situ* hybridization was performed on embryos expressing an N-terminal EGFP-tagged GAF allele independently generated through CRISPR/Cas9 genome editing. Maternal EGFP-GAF was knocked down using deGradFP expressed via the Gal4-UAS system. Gal4 expression was driven by the second chromosome maternal alpha-tubulin Gal4-VP16 driver (*mat-alpha 4-Gal4-VP16*) “64” from Bloomington Drosophila Stock Center # 80361. DegradFP expression was through *UASP-NSlmb-vhhGFP4* “2” from Bloomington Drosophila Stock Center # 38422. Embryos from *GAF^SΔPQ^/Df(3L)ED4543* (BDSC 8073) females were used to determine if GAF^SΔPQ^ can repress AAGAG RNA. Staged embryos were collected for *in situ* hybridization essentially as described above.

#### Probe Synthesis

The probe was synthesized as described in Mills et. al 2019. Template sequence listed below.

AAGAG(n)-T3as: 5’GAGAAGAGAAGAGAAGAGAAGAGAAGAGAAGAGAATCTCCCTTTAGTGAGGGTT AATT-3’

#### Immunostaining

Fixed embryos were washed 3x 10 minutes in PTx, then blocked for 1 hour at room temp. Embryos were incubated in primary antibody solution (1:1000 rabbit anti-GFP or 1:1000 rabbit anti-RNA Pol II CTD repeat YSPTSPS (phospho S5) in blocking buffer (1:10 Western Blocking Reagent (Sigma 11921673001) in PTx)) overnight at 4C, then washed 3x 10 minutes in PTx. The secondary antibody incubation (1:500 goat anti-rabbit 546 in blocking buffer) occurred for 2 hours at room temperature. Embryos were then washed 3x 10 minutes in PTx, post-fixed in a 4% paraformaldehyde solution for 20 minutes,

#### Hybridization

Fixed embryos were incubated in permeabilization solution (0.1% Triton X-100, 0.05% Igepal CA-630 (v/v), 500 ug/mL sodium deoxycholate, 500 ug/mL saponin, 2 mg/mL BSA Fraction V) for 2 hours at 4°C. Embryos were then post-fixed in a 4% paraformaldehyde solution for 20 minutes, washed 5x for 5 minutes in PTx (1x PBS, 0.1% Triton X-100), and incubated in a 1:1 Hybridization Buffer (50% deionized formamide (v/v), 25% 20x SSC (v/v), 50x) : PTx mix for 10 minutes. Prior to addition of the FISH probe, embryos were incubated in hybridization buffer for 1 hour at 55°C. A 1 ng/ul probe solution was prepared in hybridization buffer and probe hybridization occurred overnight at 55C. Post-hybridization, embryos were rinsed once in hybridization buffer, then incubated in fresh hybridization buffer for 1 hour at 55°C. Embryos were once again incubated in a 1:1 Hybridization Buffer: PTx mix for 10 minutes, followed by 5x 5 minute washes in PTx. To detect the AAGAG probe, an anti-digoxigenin Alexa 488 conjugate (Jackson ImmunoResearch 200-541-156) was diluted to 2.5 ug/mL in blocking buffer (1:10 Western Blocking Reagent (Sigma 11921673001) in PTx). Embryos were blocked for 1 hour, then incubated in antibody solution for 2 hours at room temperature. Final washes performed 3x 10 minutes in PTw (1x PBS, 0.1% Tween-20).

#### RNA-FISH embryo Imaging

Embryos were mounted in Prolong Gold Antifade Mountant (Invitrogen P10144) and imaged using Leica SP8 WLL confocal microscope. Images captured using 63x 1.3NA glycerol objective at 1024 x 512 pixels, 400 Hz scan rate. Alexa 488 and 546 were excited at 499 nm and 560 nm wavelengths respectively. Surface view images were collected with 0.30 um z-steps and max projected over a 15 μm range. RNA Pol II images captured under the same conditions described here, but with a 200 Hz scan speed and 3.0x zoom. Images shown at a single z slice.

### Assay for transposase-accessible chromatin (ATAC-seq)

Embryos from *GAF^SΔPQ^*/*Trl^s2325^* and *+*/*Trl^s2325^* females were collected from a half hour lay and aged for 2 hr. Embryos were dechorionated in bleach, mounted in halocarbon 27 oil, and hand sorted based on stage 5 morphology. Four replicates were analyzed for each genotype. Single-embryo ATAC-seq was performed as described previously (Blythe and Wieschaus 2016; Buenrostro et al. 2013). Briefly, a single dechorionated embryo was transferred to the detached cap of a 1.5 ml microcentrifuge tube containing 10 µl of ice-cold ATAC lysis buffer (10 mM Tris pH 7.5, 10 mM NaCl, 3 mM MgCl2, 0.1% NP-40). Under a dissecting microscope, a microcapillary tube was used to homogenize the embryo. The cap was placed into a 1.5 ml microcentrifuge tube containing an additional 40 µl of cold lysis buffer. Tubes were centrifuged for 10 min at 500 g at 4°C. The supernatant was removed, and the resulting nuclear pellet was resuspended in 5 µl buffer TD (Illumina, San Diego, CA) and combined with 2.5 µl H2O and 2.5 µl Tn5 transposase (Tagment DNA Enzyme, Illumina). Tubes were placed at 37°C for 30 min and the resulting fragmented DNA was purified using the Minelute Cleanup Kit (Qiagen, Hilden, Germany), with elution performed in 10 µl of the provided elution buffer. Libraries were amplified for 12 PCR cycles with unique dual index primers using the NEBNext Hi-Fi 2X PCR Master Mix (New England Biolabs). Amplified libraries were purified using a 1.2X ratio of Axygen magnetic beads (Corning Inc, Corning, NY). Libraries were submitted to the University of Wisconsin-Madison Biotechnology Center for 150 bp, paired-end sequencing on the Illumina NovaSeq 6000.

### ATAC-seq analysis

Adapter sequences were removed from raw sequence reads using NGMerge (Gaspar, 2018). ATAC-seq reads were aligned to the *Drosophila melanogaster* (dm6) genome using bowtie2 with the following parameters: -`-very-sensitive, --no-mixed, -- no-discordant, -X 5000, -k 2`. Reads with a mapping quality score < 30 were discarded, as were reads aligning to scaffolds or the mitochondrial genome. Analysis was restricted to fragments < 100 bp, which, as described previously, are most likely to originate from nucleosome-free regions (Buenrostro et al. 2013). To maximize the sensitivity of peak calling, reads from all replicates of *GAF^SΔPQ^* and control embryos were combined. Peak calling was performed on combined reads using MACS2 with parameters `-f BAMPE --keep-dup all -g 1.2e8 --call-narrowPeak`. Reads aligning within accessible regions were quantified using featureCounts, and differential accessibility analysis was performed using DESeq2 with an adjusted p-value<0.05 and a fold change > 2 as thresholds for differential accessibility. To determine overlap of accessible peaks with GAF ChIP-seq data, we used intersectBed from Bedtools to compare accessible peaks to high confidence peaks from stage 5 sfGFP-GAF(N) ChIP-seq (Gaskill et al. 2021). To determine overlap of accessible peaks with peaks called as differential in GAF^deGradFP^ embryos, we used intersectBed from Bedtools to compare accessible peaks to all differential peaks from GAF^deGradFP^ ATAC-seq (Gaskill et al. 2021).

Peaks with at least a 5% overlap were considered to be shared. The volcano plot was generated with ggplot. Genome browser tracks were generated using z-score normalized bigWigs with IGV. Numbers used for all Fisher’s exact tests are included in Table S3. Pearson’s correlation coefficients were calculated for all ATAC-seq replicates using deeptools (Ramírez et al. 2016) and are reported in Table S3.

## Data availability

Sequencing data have been deposited in GEO under accession code GSE218020.

## Acknowledgements

We thank Gary Karpen, Bloomington Drosophila Stock Center, and the Drosophila Genome Resource Center for providing reagents and fly lines. We acknowledge the University of Wisconsin-Madison Biochemistry Department Optical Core for access to microscopes for imaging and the University of Wisconsin-Madison Biotechnology Center and the NUSeq Core Facility for sequencing. Elizabeth Larson assisted with trouble-shooting the data analysis. MMG and TJG were supported by National Institutes of Health (NIH) National Research Service Award T32 GM007215. APB was supported by NIH National Research Service Award T32 GM132039. Experiments were supported by a R35 GM136298 from NIH (MMH), NIH DP2OD020635 (MM), NIH R01HD101563 (SAB). MMH was also supported by a Vallee Scholar Award (MMH). MM was also supported by a grant from Margaret Q Landenberger Foundation. SAB is a Pew Scholar in the Biomedical Sciences, supported by The Pew Charitable Trusts.

## Supplemental Figure Legends

**Figure S1: GAF forms multiple, stable nuclear foci during the MZT.** A. Quantification of the percent of the total volume of the nucleus occupied by sfGFP-tagged GAF foci (*top*), sfGFP-tagged GAF foci volume (*middle*), and number of sfGFP-tagged GAF foci per nucleus (*bottom*) during NC13 and NC14. B. Quantification of the volume of sfGFP-tagged GAF foci in NC13. C. Quantification of the number of sfGFP-tagged GAF foci per nucleus during NC13. Asterisks indicate pairwise p-value thresholds (** = 0.01, *** = 0.001, **** = 0.0001) calculated with Tukey-Kramer test. n = 2 embryos, 14 nuclei analyzed. D. Representative lattice light-sheet images of sfGFP-tagged GAF foci during NC13 and NC14. Scale bars, 2.5 μm

**Figure S2: Neither the long isoform nor the short isoform polyQ domain is required for viability.** A. Cartoon representation of the protein domains of the two GAF isoforms. B. Sashimi plots generated from modENCODE RNA-seq data showing the differential splice junctions used for the two GAF isoforms in 0-2 hr AEL embryos, 2-4 hr AEL embryos, and 22-24 AEL embryos. The black arrowhead identifies the splice junctions used for the long GAF isoform. C. PCR products amplified from cDNA extracted from *GAF^L^*, *GAF^SΔPQ^,* and *w^1118^* overnight embryos with primers F1 and R1 as indicated in B. The expected sizes of the products are wild-type long isoform = 1522 bp; wild-type short isoform = 2034 bp; truncated short isoform = 1737 bp. Isoform-specific products are marked by colored arrowheads, and those verified by sequencing are indicated with an asterisk. D. PCR products amplified from cDNA extracted from *GAF^L^*, *GAF^SΔPQ^,* and *w^1118^* overnight embryos with primers F2 and R2 as indicated in B. The expected sizes of the products are wild-type short isoform = 1310 bp; truncated short isoform = 1013 bp. E. Western blot with anti-GAF antibody on embryo extracts from *GAF^L^/-*, *GAF^SΔPQ^/-,* and *+/-* overnight embryos. The short isoform is detected (green arrow) and truncated product is evident in *GAF^SΔPQ^/-* embryos (blue arrow). The long isoform is not clearly detected. F. Percent of balancer (TM3) to nonbalancer adults for the crosses resulting in the indicated nonbalancer progeny. Heterozygous parents were mated, and progeny were scored for the presence of the balancer. n = total number of flies assayed. G. Western blot with anti-GFP antibody on embryo extracts from *sfGFP-GAF^SΔPQ^/+, sfGFP-GAF/+,* and *w^1118^* overnight embryos. The tagged GAF protein is detected (green arrow), and the truncated tagged product is evident in *sfGFP-GAF^SΔPQ^/+* embryos (blue arrow). Tubulin is a loading control.

**Figure S3: GAF DNA binding is required for localization to foci and mitotic retention.** A-B. Images of NC14 embryos expressing sfGFP-tagged GAF transgenes that localize to foci (A) or do not localize to foci (B). Embryos were laid by mothers expressing endogenous mCherry-GAF or His2Av-RFP and the sfGFP-tagged GAF transgene as indicated. Anaphase images were generated from embryos expressing sfGFP-tagged GAF transgenes and His2Av-RFP to mark the mitotic chromosomes. mCherry-GAF or His2Av-RFP is in magenta. sfGFP-tagged GAF is in green. FL = Full-length. Scale bars 5μm.

**Figure S4: GAF is localized to subnuclear foci that are distinct from many other proteins.** Interphase NC14 embryos laid by mothers expressing endogenous mCherry-GAF and the foci forming GFP-tagged factor indicated above. mCherry-GAF is in magenta. The GFP-tagged factors are in green. Closed arrows indicate mCherry-GAF only foci. Open arrows indicate foci formed by GFP-tagged proteins alone. The arrowhead indicates a region of overlap. Scale bars 5μm.

**Figure S5: GAF is not required for global 3D chromatin organization.** A. Hi-C contact map of Chr 3R 14.4-15.2 Mb for GAF^deGradFP^ and control (*sfGFP-GAF(N)* homozygous) embryos collected at 2-2.5 hr AEL (*top*). ChIP-seq signal from stage 5 GAF-sfGFP embryos (Gaskill et al. 2021) and the insulation score for GAF^deGradFP^ embryos and paired controls over the same genomic region (*bottom*). B. Compartment scores over Chr. 2L for GAF^deGradFP^ and control embryos. C. Hi-C contact maps for the *Antennapedia* (*Antp*) locus for control (*top*) and GAF^deGradFP^ (*bottom*). Stage 5 GAF-sfGFP ChIP-seq signal (Gaskill et al. 2021) is shown on the right. The red circle highlights a loop between the *Antp* promoter and an upstream GAF-bound region that is lost in the GAF^deGradFP^ embryos. D. DNA-FISH on GAF^deGradFP^ embryos at NC14 using an (AAGAG)7 probe. Anti-GFP immunostaining confirms the absence of sfGFP-GAF. Scale bars, 5 μm.

**Figure S6: RNA Pol II co-localizes with AAGAG transcripts in GAF^deGradFP^ embryos.** AAGAG RNA FISH with RNA Pol II immunostaining performed in EGFP-GAF and GAF^deGradFP^ NC14 embryos. The intensity of the AAGAG RNA channel in EGFP-GAF embryos was scaled to highlight foci. The circle indicates a representative nucleus showing AAGAG RNA co-localized with a high-density region of RNA Pol II. Scale bars are 5 μm.

**Supplemental Table S1: Peaks called in ChIP-seq for 2-2.5 hr AEL, hand-sorted embryos laid by *sfGFP-GAF/+* and *sfGFP-GAF^SΔPQ^/+* females.** Chromosome, start and end for each peak are provided as labelled.

**Supplemental Table S2: Differential peaks identified in ATAC-seq of 2-2.5 hr AEL, hand-sorted embryos laid by *GAF^SΔPQ^/-* females compared to *+/-* controls.** Data are provided in the first sheet and columns are defined in the second sheet.

**Supplemental Table S3: Numbers for statistical analyses performed.**

**Supplemental Movie S1: Video acquired using lattice light-sheet microscopy of a NC14 embryo expressing HP1a-RFP and sfGFP-tagged GAF.** sfGFP-tagged GAF is in green. HP1a-RFP is in magenta.

**Supplemental Movie S2: Video acquired using lattice light-sheet microscopy of an embryo throughout NC11-NC14 expressing HP1a-RFP and sfGFP-tagged GAF.** sfGFP-tagged GAF is in green. HP1a-RFP is in magenta.

