## Supplemental Figures for "Localization of the pioneer factor GAF to subnuclear foci is driven by DNA binding and required to silence satellite repeat expression"

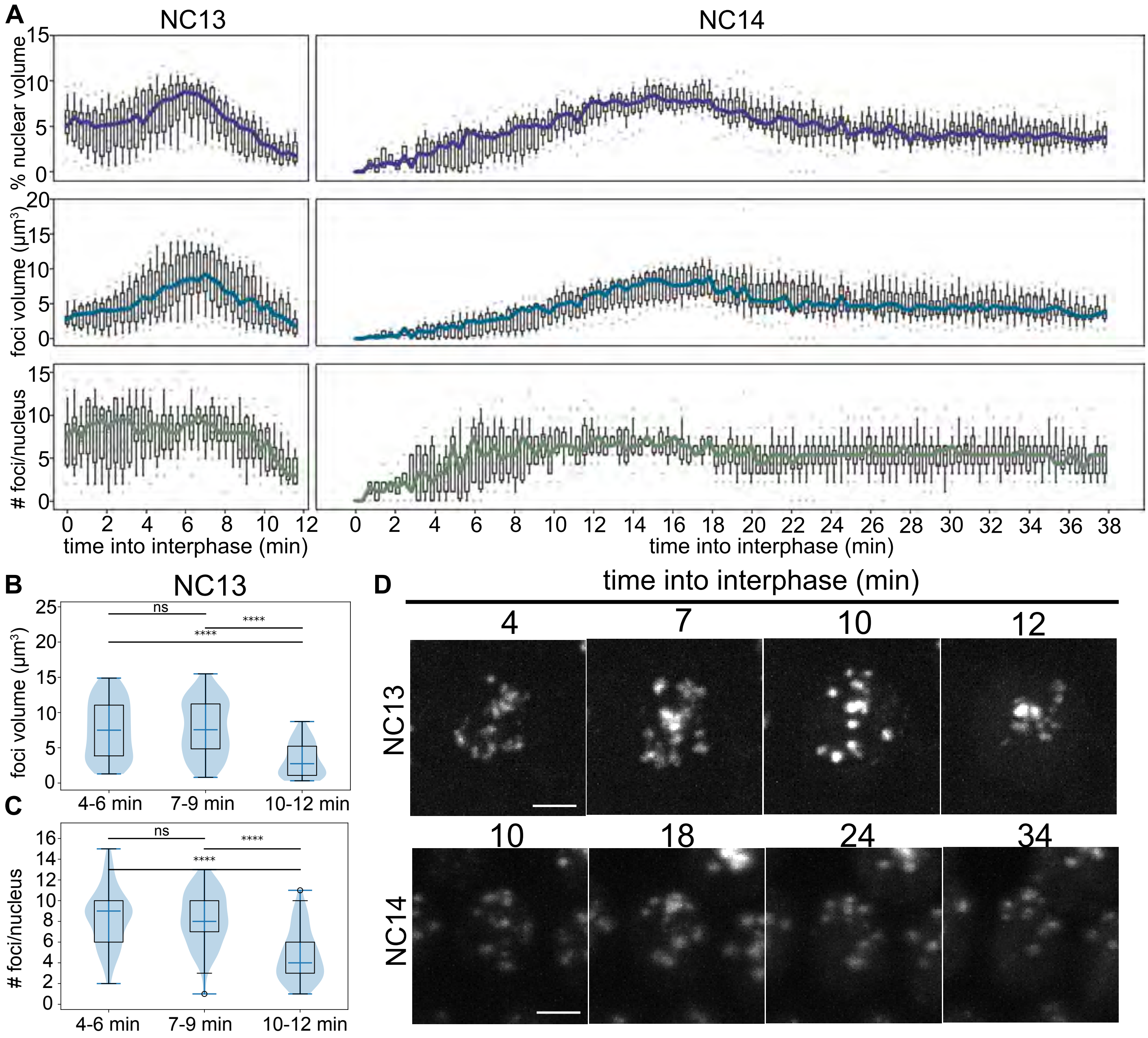


**Figure S1: GAF forms multiple, stable nuclear foci during the MZT.** A. Quantification of the percent of the total volume of the nucleus occupied by sfGFP-tagged GAF foci (*top*), the sfGFP-tagged GAF foci volume (*middle*), and number of sfGFP-tagged GAF foci per nucleus (*bottom*) during NC13 and NC14. B. Quantification of the volume of sfGFP-tagged GAF foci in NC13. C. Quantification of the number of sfGFP-tagged GAF foci per nucleus in NC13. Asterisks indicate pairwise p-value thresholds (** = 0.01, *** = 0.001, **** = 0.0001) calculated with the Tukey-Kramer test. n = 2 embryos, 14 nuclei analyzed. D. Representative lattice light-sheet images of sfGFP-tagged GAF foci during NC13 and NC14. Scale bar, 2.5 $\mu$m.

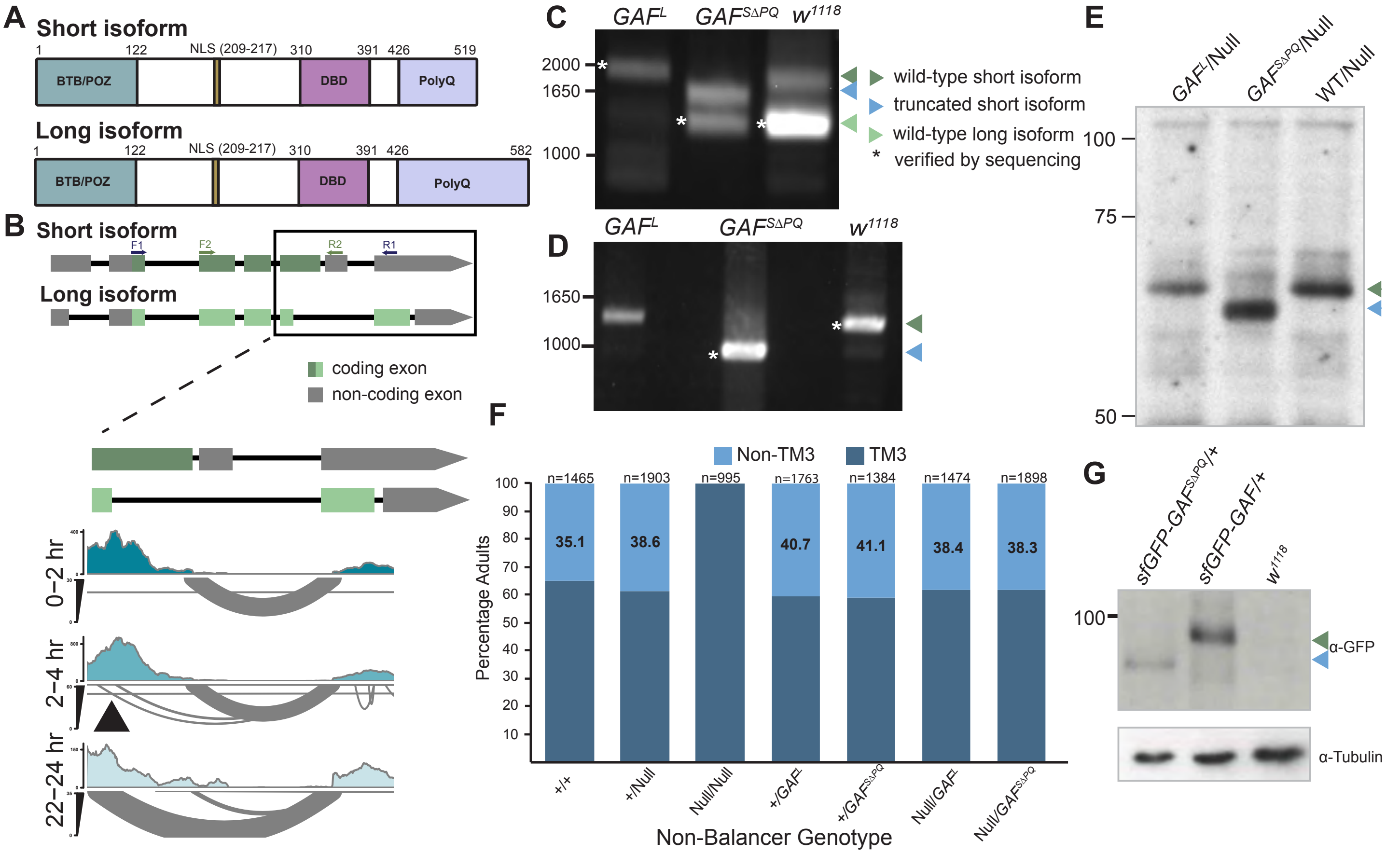


**Figure S2: Neither the long isoform nor the short isoform polyQ domain is required for viability**. A. Cartoon representation of the protein domains of the two GAF isoforms. B. Sashimi plots generated from modENCODE RNA-seq data showing the differential splice junctions used for the two GAF isoforms at 0-2 hr AEL embryos, 2-4 hr AEL embryos, and 22-24 hr AEL embryos. The black arrowhead identifies the splice junctions used for the long GAF isoform. C. PCR products amplified from cDNA extracted from *GAF$^{L}$*, *GAF$^{S\Delta PQ}$*, and *w$^{1118}$* overnight embryos with primers F1 and R1 as indicated in B. The expected sizes of the products are: wild-type long isoform, 1522 bp; wild-type short isoform, 2034 bp; truncated short isoform,1737 bp. Isoform-specific products are marked by colored arrowheads, and those verified by sequencing are indicated with an asterisk. D. PCR products amplified from cDNA extracted from *GAF$^{L}$*, *GAF$^{S\Delta PQ}$*, and *w$^{1118}$* overnight embryos with primers F2 and R2 as indicated in B. The expected sizes of the products are: wild-type short isoform, 1310 bp; truncated short isoform,1013 bp. E. Western blot with anti-GAF antibody on embryo extracts from *GAF$^{L}$*/-, *GAF$^{S\Delta PQ}$*/-, and +/- overnight embryos. The short isoform is detected (green arrow), and a truncated product is evident in *GAF$^{S\Delta PQ}$*/- embryos (blue arrow). The long isoform is not clearly detected. F. Percent of balancer (TM3) to nonbalancer adults for the crosses resulting in the indicated nonbalancer progeny. Heterozygous parents were mated, and progeny were scored for the presence of the balancer. n = total number of flies assayed. G. Western blot with anti-GFP antibody on embryo extracts from *sfGFP-GAF$^{S\Delta PQ}$/+*, *sfGFP-GAF/+*, and *w$^{1118}$* overnight embryos. The tagged GAF protein is detected (green arrow), and the truncated-tagged product is evident in *sfGFP-GAF$^{S\Delta PQ}$/+* embryos (blue arrows). Tubulin is a loading control.

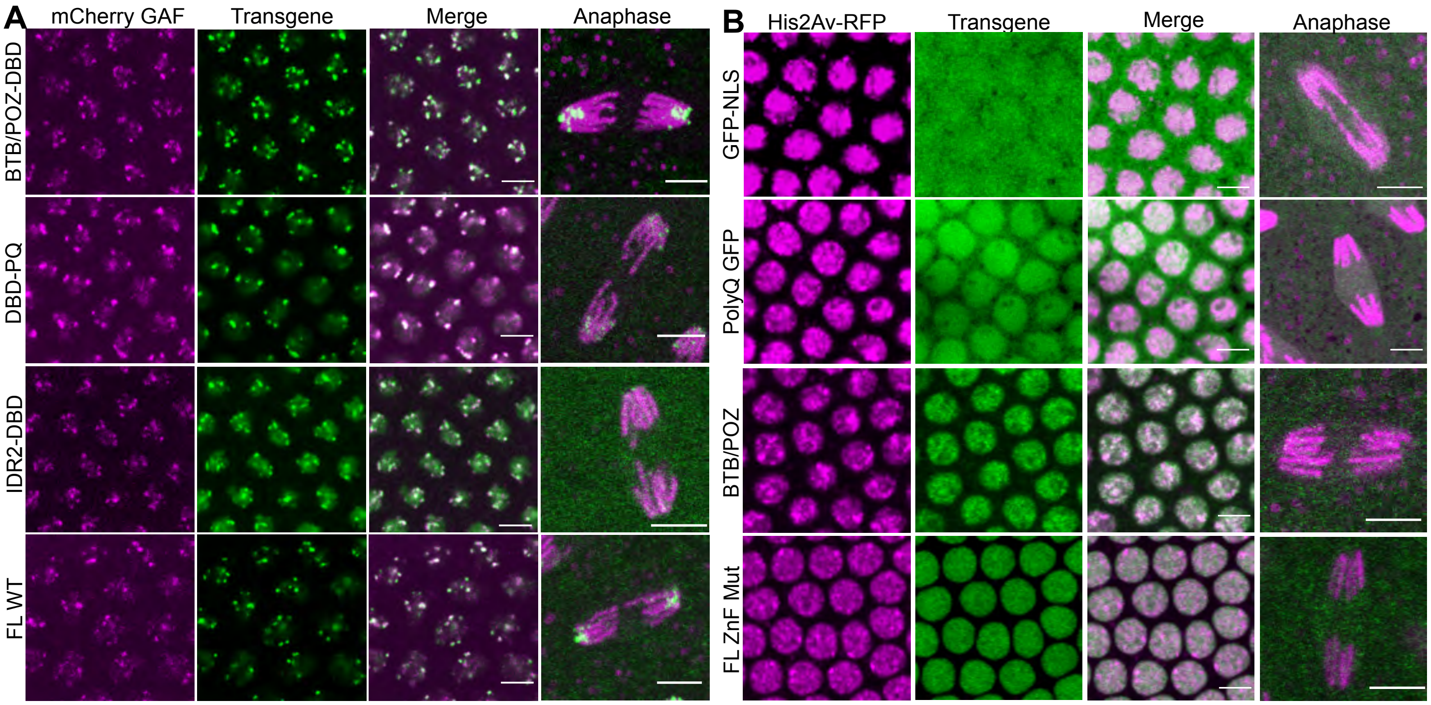


**Figure S3 : GAF DNA-binding is required for localization to foci and mitotic retention.** A-B. Images of NC14 embryos expressing sfGFP-GAF transgenes that localize to foci (A) or do not localize to foci (B). Embryos were laid by mothers expressing endogenous mCherry-GAF or His2Av-RFP and the sfGFP-tagged GAF transgene as indicated. Anaphase images were generated from embryos expressing sfGFP-tagged GAF transgenes and His2Av-RFP to mark ths chromosomes. mCherry-GAF or His2Av-RFP is in magenta. sfGFP-GAF is in green. FL = full length Scale bars are 5μm.

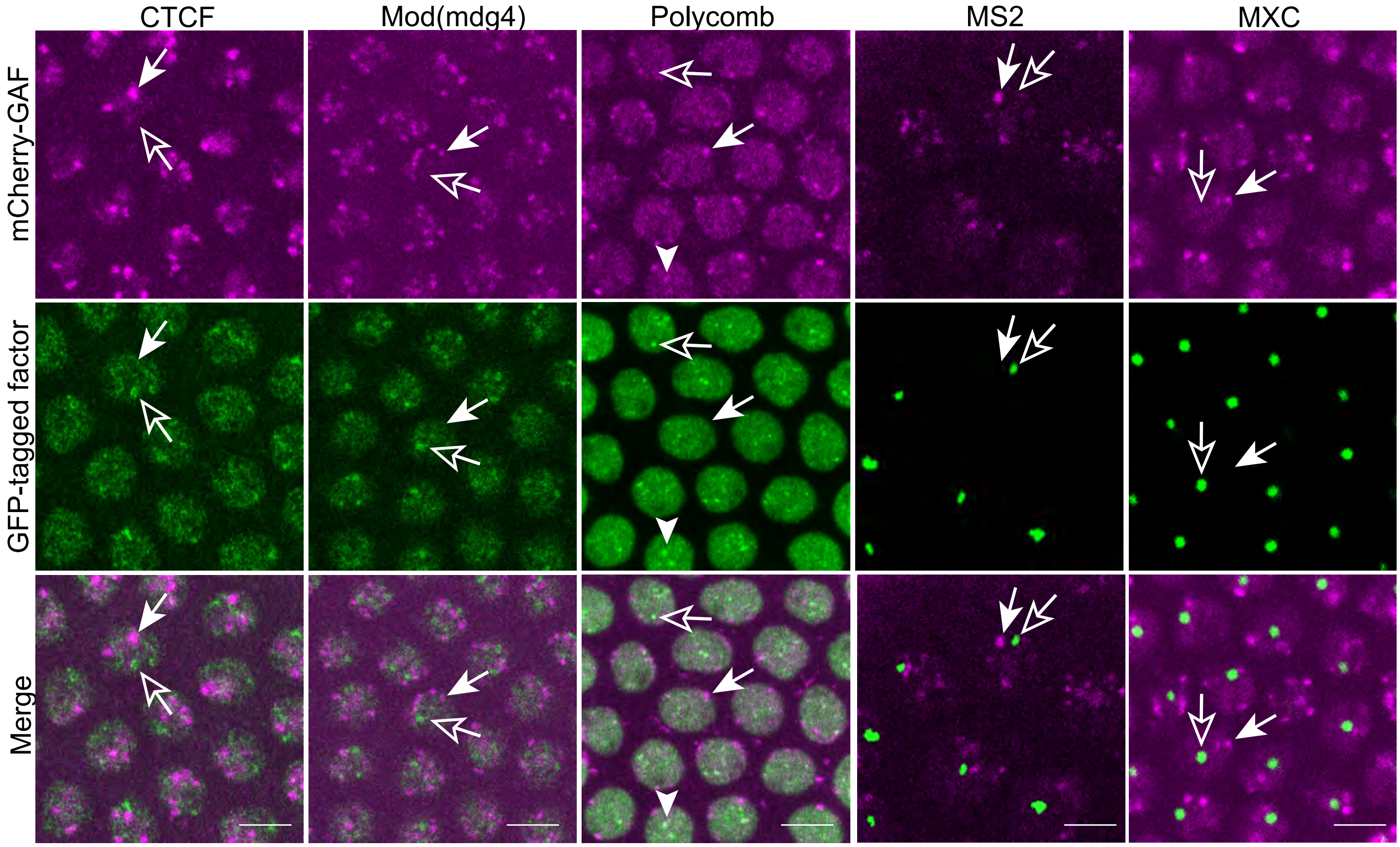


**Figure S4: GAF is localized to subnuclear foci that are distinct from many other proteins.** Interphase NC14 embryos laid by mothers expressing endogenous mCherry-GAF and the foci forming GFP-tagged factor indicated above. mCherry is in magenta. The GFP-tagged factors are in green. Closed arrows indicate mCherry-GAF only foci. Open arrows indicate foci formed by GFP-tagged proteins alone. The arrowhead indicates a region of overlap.Scale bars 5µm.

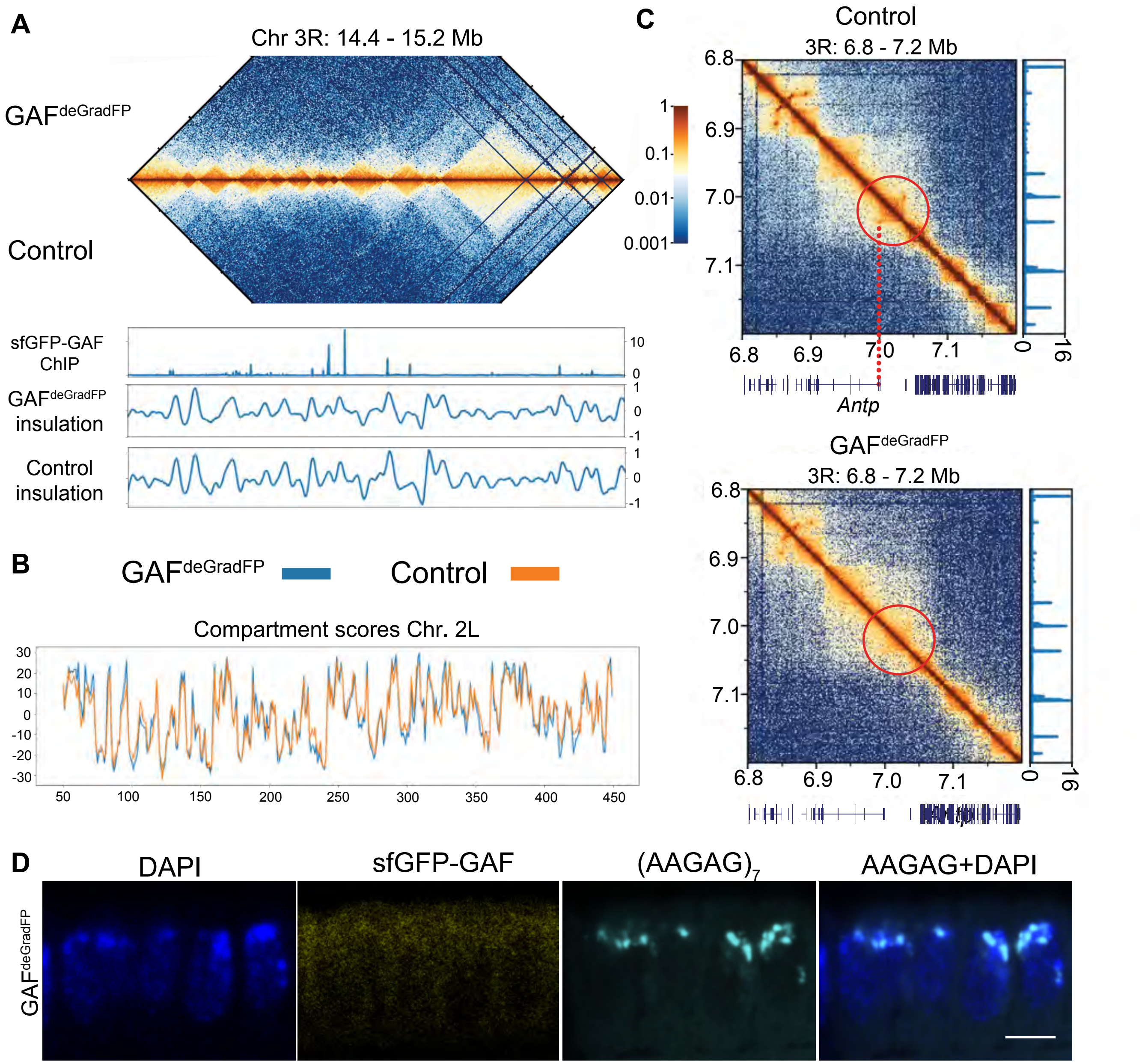


**Figure S5: GAF is not required for global 3D chromatin organization**. A. Hi-C contact map of Chr 3R: 14.4-15.2 Mb for GAF$^{deGradFP}$ and control (*sfGFP-GAF(N)* homozygous) embryos collected at 2-2.5hr AEL (*top*). ChIP-seq signal from stage 5 sfGFP-GAF embryos (Gaskill et al. 2021) and the insulation scores for GAF$^{deGradFP}$ embryos and paired controls over the same genomic region (*bottom*). B. Compartment scores over Chr. 2L for GAF$^{deGradFP}$ and control embryos. C. Hi-C contact maps for the *Antennapedia* (*Antp*) locus for control (*top*) and GAF$^{deGradFP}$ (*bottom*). Stage 5 GAF-sfGFP ChIP-seq signal (Gaskill et al. 2021) is shown on the right. The red circle highlights a loop between the *Antp* promoter and an upstream GAF-bound region that is lost in the GAF$^{deGradFP}$ embryos. D. DNA-FISH on GAF$^{deGradFP}$ embryos at NC14 using an $(AAGAG)_7$ probe. Anti-GFP immunostaining demonstrates the absence of sfGFP-GAF. Scale bars, 5µm.

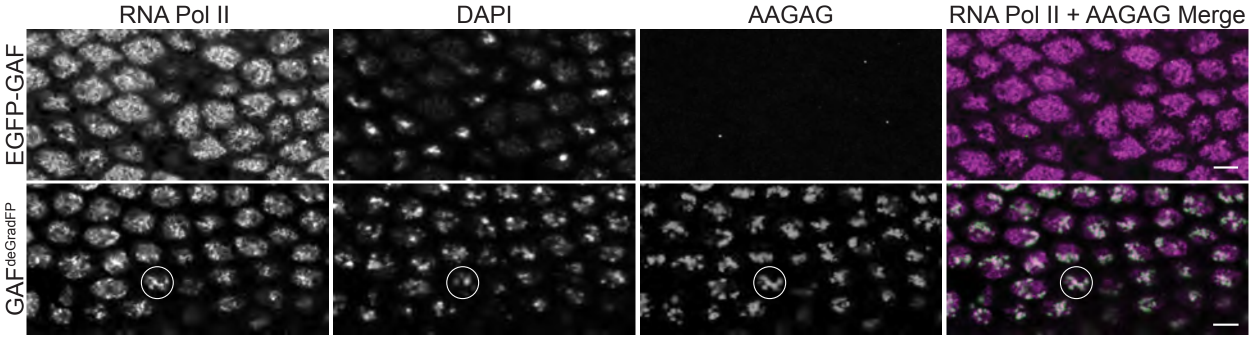


**Figure S6: RNA Pol II co-localizes with AAGAG transcripts in GAF$^{deGradFP}$ embryos.** AAGAG RNA FISH with RNA Pol II immunostaining performed in EGFP-GAF and GAF$^{deGradFP}$ NC14 embryos. The intensity of the AAGAG RNA channel in EGFP-GAF embryos was scaled to highlight foci. The circle indicates a representative nucleus showing AAGAG RNA co-localized with a high-density region of RNA Pol II. Scale bars are 5 μM.
